# Selective targeting of non-centrosomal AURKA functions through use of a targeted protein degradation tool

**DOI:** 10.1101/2020.07.22.215814

**Authors:** Richard Wang, Camilla Ascanelli, Ahmed Abdelbaki, Alex Fung, Tim Rasmusson, Iacovos Michaelides, Karen Roberts, Catherine Lindon

## Abstract

Targeted protein degradation tools are becoming a new therapeutic modality, allowing small molecule ligands to be reformulated as heterobifunctional molecules (PROteolysis Targeting Chimeras, PROTACs) that recruit ubiquitin ligases to targets of interest, leading to ubiquitination and destruction of the targets. Several PROTACs against targets of clinical interest have been described, but detailed descriptions of the cell biology modulated by PROTACs are missing from the literature. Here we describe the functional characterization of a PROTAC derived from AURKA inhibitor MLN8237 (alisertib). We demonstrate efficient and specific destruction of both endogenous and overexpressed AURKA by Cereblon-directed PROTACs. At the subcellular level, we find differential targeting of AURKA on the mitotic spindle compared to centrosomes. The phenotypic consequences of PROTAC treatment are therefore distinct from those mediated by alisertib, and in mitotic cells differentially regulate centrosome- and chromatin-based microtubule spindle assembly pathways. In interphase cells PROTAC-mediated clearance of non-centrosomal AURKA modulates the cytoplasmic role played by AURKA in mitochondrial dynamics, whilst the centrosomal pool is refractory to PROTAC-mediated clearance. Our results point to differential sensitivity of subcellular pools of substrate, governed by substrate conformation or localization-dependent accessibility to PROTAC action, a phenomenon not previously described for this new class of degrader compounds.

## Introduction

The advent of targeted protein degradation tools that exploit the endogenous protein degradation machinery to eliminate disease proteins from the cell has started a revolution in therapeutic strategy and drug design^1^. One novel way to target proteins for degradation is through PROteolysis Targeting Chimeras (PROTACs), consisting of a chimeric molecule that binds at one end to a protein target, and at the other to a ubiquitin ligase (E3), most commonly the Cereblon (CRBN) substrate recognition protein together with the CUL4A E3 ligase complex, or to the von Hippel Lindau (VHL) protein in association with the CUL2 complex^2,3^. This PROTAC-mediated ternary complex formation between functional E3 and target protein facilitates ubiquitin transfer, leading to ubiquitination of the target and its proteolysis at the 26S proteasome.

This new paradigm of ‘event-driven’ pharmacology (in contrast to the use of traditional ‘occupancy-based’ drugs) holds great hope for the development of catalytic drugs able to work at lower doses and with higher specificity than the ligands from which they are derived. Moreover, the altered pharmacodynamics of substrate destruction versus inhibition raises the possibility of repurposing small molecule ligands (including those that have failed clinical trials as inhibitors of their targets) into PROTACs. However, although a number of publications document the success of novel PROTACs in eliminating their cellular targets, there has been little impact of this technology so far in the field of cell biology. PROTACs have clear potential as tools for cell biology that can perturb cellular protein functions on a timescale more favourable than siRNA-mediated interference and in a way that does not depend on effective inhibition of an enzymatic function. In this study, we investigate the properties of a PROTAC tool based on a known small molecule inhibitor of the mitotic kinase Aurora A (AURKA), MLN8237 (also known as alisertib)^4–6^.

AURKA is a well-studied regulator of mitosis, playing critical roles in centrosome maturation, mitotic timing, microtubule nucleation and spindle assembly^7,8^. Distinct populations of AURKA are either recruited to centrosomes by CEP192, or on spindle microtubules (MTs) via the MT-associated protein TPX2. These separate populations can be independently perturbed through disruption of either interaction^9–11^. AURKA activity at centrosomes contributes to mitotic entry. Activation of AURKA is thought to occur either through auto-phosphorylation in the T-loop (at T287/288), a process promoted by CEP192 oligomerization at the centrosomes, or through interaction with a number of known binding partners that act to stabilize the ‘DFG-In’ conformation to favour kinase activity independently of T-loop phosphorylation^12–14^. The best-known of these interactors is TPX2. At nuclear envelope breakdown (NEB), TPX2 is released by importin-α, under the influence of the RanGTP gradient around the mitotic chromosomes, to bind and activate AURKA. *In vitro* tests show that binding by TPX2 and T-loop phosphorylation independently activate AURKA approximately 100-fold^15,16^. These separable intracellular AURKA activities (defined by pT288 at the centrosomes and TPX2 binding around chromatin) contribute to distinct pathways of MT nucleation that act together to achieve mitotic spindle assembly. Critical targets of AURKA in both pathways are NEDD1 and TACC3. Recruitment and phosphorylation of NEDD1 allows recruitment of the γ-TURC nucleating complex whilst phosphorylation of TACC3 promotes assembly of a pTACC3-ch-TOG-clathrin complex^17^ proposed to stabilize parallel MTs in the spindle.

AURKA undergoes targeted proteolysis in every cell cycle as a substrate of the Anaphase-Promoting Complex (APC/C) ubiquitin ligase at mitotic exit^18,19^. However, AURKA is detectable in interphase cells and has been attributed a number of non-mitotic roles including ciliation control, cell cycle regulation of MYCN-dependent transcription, DNA damage pathways and mitochondrial regulation^20–23^. Overall, there is a growing interest in the roles played by AURKA outside of mitosis and their contribution to its cancer-promoting activity. AURKA has long been a postulated therapeutic target due to its well-documented overexpression in cancer, although the role it plays in oncogenesis is far from clear. Recent structural and conformational studies have led to improved understanding of its mode of activation and the realization that multiple active forms may persist through interphase that depend on different binding partners. Recent work from our lab has shown that un-degraded AURKA retains activity after mitosis^24^.

Therefore, a PROTAC tool able to eliminate AURKA protein could be an important cell biology tool as well as a potential therapeutic strategy. Here we test characteristics of PROTAC activity directed against AURKA and investigate the cell biology that accompanies targeted protein degradation of this critical cellular target.

## Results

### AURKA-Venus is sensitive to CRBN-directed PROTACs

We set out to investigate the action of AURKA-directed targeted protein degradation tools (PROTACs) against AURKA in single cell time-lapse assays using cell lines that we have previously described^25^: an AURKA-Venus knock-in line in RPE1 cells (AURKA-Venus^KI^) where AURKA-Venus recapitulates expression of the endogenous protein (undetectable in interphase cells and strongly upregulated for mitosis), and a line expressing exogenous AURKA-Venus under tetracycline control (RPE1FRT/TO-AURKA-Venus, AURKA-Venus^TO^) where higher levels of expression occur throughout the cell cycle. We used AURKA-Venus^KI^ and AURKA-Venus^TO^ cells arrested in mitosis by an agonist of the Spindle Assembly Checkpoint (SAC), STLC, to test the activity of PROTAC compounds (Figure 1).

**Figure 1.**
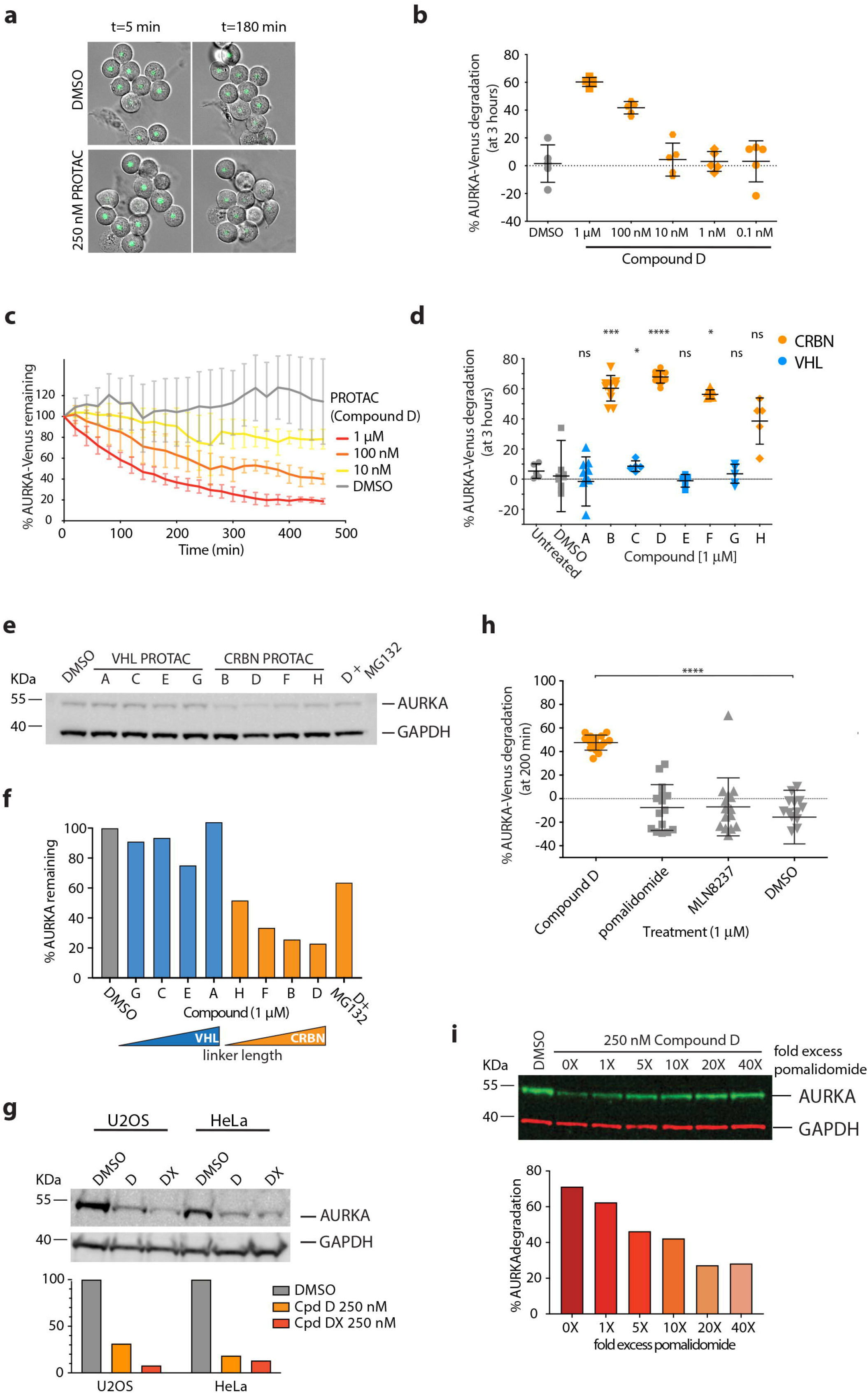
AURKA destruction following treatment with CUL4-based PROTACs. **a-d** AURKA-Venus^KI^ degradation in STLC-arrested RPE-1 cells, measured by quantitative timelapse imaging. **a** Examples of fields of cells treated with a PROTAC (Compound D) or vehicle control (DMSO). Venus fluorescence was measured in individual cells and plotted as an end-point assay (percentage of AURKA-Venus^KI^ degradation after 3 hr) (**b**) or as percentage of AURKA-Venus^KI^ remaining over time (to show the kinetics of degradation) (**c**). **b** Scatter plots showing response to different doses of PROTAC (Compound D) or DMSO, with whiskers indicating mean ± SDs. **c** Time-course of AURKA-Venus^KI^ degradation plotted as mean fluorescence ± SDs at each time point (n ≥ 7 cells). **d** Comparison of percentage degradation of AURKA-Venus^KI^ after treatment with potential PROTAC compounds directed to CUL4A (via CRBN) or CUL2 (VHL) ubiquitin ligases (listed in Table 1). Scatter plots show pooled results from two separate experimental repeats with whiskers indicating mean values and SDs. Results of Kruskal-Wallis multiple ANOVA, and the Dunn’s post-hoc multiple comparison test to DMSO are indicated. **e-f** Degradation of endogenous AURKA in HeLa cells following 3 hr treatment with PROTACs, with or without 42 µM proteasome inhibitor MG132, measured by quantitative immunoblotting (**e**) to plot percentage protein remaining, ordered to show increasing linker size across the series (**f**). **g** U2OS and HeLa cells were treated for 3 hours with 250 nM Compound D (PROTAC-D) or a new compound with increased linker length (PROTAC-DX). Endogenous AURKA levels were measured by immunoblot and plotted as percentage remaining compared to DMSO treatment after normalization to GAPDH loading control. The experiment shown is one of two repeats that gave identical results. **h** Degradation of AURKA-Venus^KI^ was measured in prometaphase cells treated with 1 μM PROTAC-D, MLN8237 or Pomalidomide. Kruskal-Wallis multiple ANOVA, and the Dunn’s post-hoc multiple comparison test to DMSO, showed that only PROTAC-D caused significant AURKA-Venus degradation. **i** Degradation of endogenous AURKA was quantified by immunoblot from mitotic-enriched U2OS cells treated with 250 nM PROTAC D in the presence of 1 – 40-fold molar excess of Pomalidomide. Data representative of two experiments (the other in HeLa cells). Only cells that did not exit mitosis during filming (scored as onset of cortical contractility followed by respreading) were included in the analyses shown in Figure 1a-d, h).

We synthesised eight PROTAC molecules consisting of the well-characterised inhibitor of AURKA, MLN8237, linked to either a known ligand of von Hippel-Lindau (VHL) E3 ligase or to the thalidomide derivative, pomalidomide, to recruit Cereblon (CRBN) E3 ligase. As the linker is an integral part of the PROTAC molecule and linker length can be a key determinant of PROTAC function, we designed four molecules for each of the MLN8237-CRBN or -VHL combinations with varying polyethylene glycol (PEG) linker lengths (Table 1).

**Table 1:** Summary of compounds tested.

| **Cpd** | **Substrate ligand**<br>**(‘warhead’)** | **E3 ligand**<br>**target** | **linker length**<br>**(M<sub>r</sub>)\*** |
| --- | --- | --- | --- |
| **A** | MLN8237 (alisertib) | VHL | 288 |
| **B** | MLN8237 | CRBN | 272 |
| **C** | MLN8237 | VHL | 200 |
| **D** | MLN8237 | CRBN | 316 |
| **DX** | MLN8237 | CRBN | 404 |
| **E** | MLN8237 | VHL | 244 |
| **F** | MLN8237 | CRBN | 228 |
| **G** | MLN8237 | VHL | 156 |
| **H** | MLN8237 | CRBN | 184 |

\*M<sub>r</sub>, relative molecular weight

We found that CRBN-based PROTAC compounds were able to elicit destruction of both AURKA-Venus^KI^ and AURKA-Venus^TO^ in time-lapse movies of mitotic arrested cells (Figure 1a-d). Compound D reduced AURKA levels in a dose dependent manner (Figure 1b, c) with an EC_50_ in the 100 nM range (Figure 1b). At a dose of 1µM, Compound D caused loss of AURKA-Venus with t_1/2_ approximately 2 hours (Figure 1c). The activity of the PROTAC against AURKA-Venus^KI^ in time-lapse assays (Figure 1d), or against endogenous AURKA in extracts from mitotic arrested HeLa cells (Figure 1e, f), appeared to correlate with linker length, suggesting that topological constraints limit the efficacy of PROTAC action. The VHL-based PROTACs tested were inactive in all but one dose (Figure 1d). Taken together, the most efficient PROTAC tested in these initial experiments was Compound D, which we named AURKA-PROTAC-D (PROTAC-D). We tested the correlation between linker length and efficacy of the PROTAC by creating a new compound with an extra-long linker, Compound DX (Table 1). As predicted, Compound DX reduced AURKA levels more efficiently than PROTAC-D (Figure 1g). We tested the specificity of PROTAC action of CRBN-directed compounds in further experiments (Figure 1h, i) demonstrating that neither MLN8237 nor the CRBN ligand (pomalidomide) on its own affected AURKA levels (Figure 1h). In addition, the action of Compound D was blocked by competition with excess pomalidomide (Figure 1i), supporting that recruitment of CRBN E3 holo-complex was necessary for AURKA level reduction.

### PROTAC-mediated degradation of AURKA-Venus is independent of mitotic pathways of destruction

While analysing these experiments, we noticed that AURKA-Venus^KI^ cells arrested in mitosis with STLC were more likely to exit mitosis after treatment with PROTAC than after treatment with DMSO. As we wanted to be able to separate PROTAC treatment effects caused by target degradation from any residual inhibitory effects caused by on-target engagement, we used Compound A (Cpd A) as a negative control. Cpd A is a MLN8237-VHL molecule of identical linker length to PROTAC-D that did not cause degradation of AURKA (Figure 1d). It had a small and non-significant effect in promoting mitotic slippage compared to PROTAC-D (Figure 2a). Since AURKA is itself a substrate of mitotic exit degradation under control of the APC/C^FZR1^, failure of the SAC, leading to activation of the APC/C, would be predicted to result in degradation of AURKA independently of PROTAC-mediated ubiquitination. Therefore, in the single cell mitotic degradation assays shown in Figure 1, we quantified only cells that remained arrested in mitosis for the duration of the assay. However, we also carried out experiments to test directly whether mitotic degradation pathways were involved in PROTAC-D-driven disappearance of AURKA-Venus, using a combination of drugs (APCin, proTAME) that inhibits the activity of the APC/C ubiquitin ligase^26^. Degradation of AURKA-Venus in response to PROTAC-D was not prevented by inhibition of APC/C (Figure 2b, c) and was therefore independent of mitotic exit.

**Figure 2.**
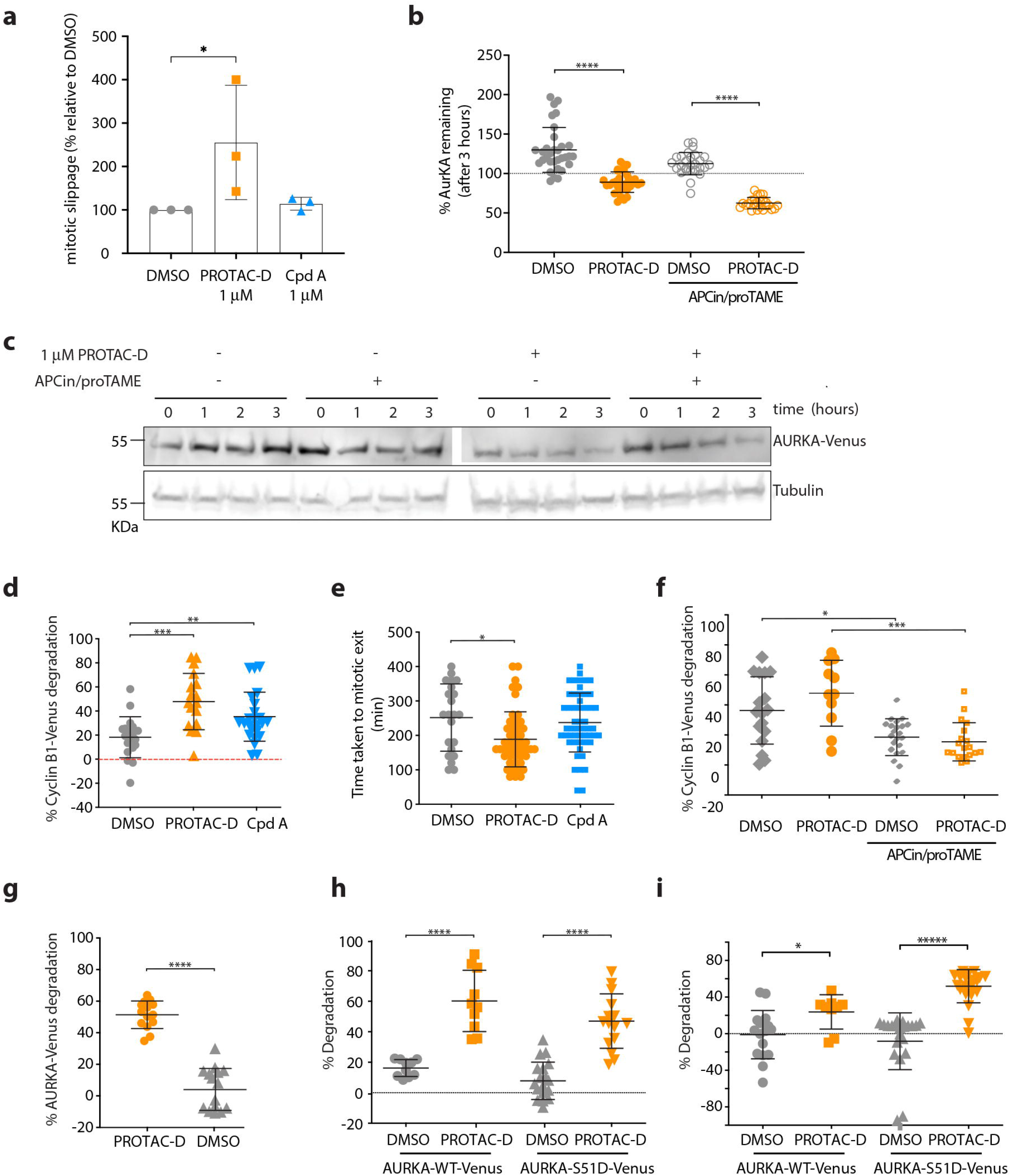
CUL4-mediated AURKA destruction is independent of known pathways of AURKA degradation. **a** RPE1-AURKA-Venus^KI^ cells treated as in Figure 1D were scored for mitotic slippage (exit from mitosis in presence of STLC). Percentage of cells undergoing slippage after 3 hr treatment with PROTAC-D or Cpd A is shown relative to DMSO-treated cells. Mean values were generated from 3 identical experiments (where n ≥ 30 cells), plotted with error bars to indicate SDs and tested by ordinary one-way ANOVA with Dunnett’s post-hoc test for significance. **b-c** Degradation of AURKA-Venus in mitotic-arrested RPE1-AURKA-Venus^KI^ cells after treatment with PROTAC-D in the presence of APCin/proTAME. Degradation was measured in single cell assays, with data shown as scatter plots (bars to indicate means ± SDs) tested with unpaired *t*-Test or Mann Whitney *U*-test for significance (**b**) or visualised as a time-course of drug treatment by immunoblot (**c**). **d-f** RPE1-Cyclin B1-Venus^KI^ cells treated with PROTACs to investigate mitotic slippage. Increased degradation of Cyclin B1–Venus following PROTAC-D treatment (**d**), correlates with increased rate of mitotic exit (**e**) (both results statistically tested by ordinary one-way ANOVA with Dunnett’s post-hoc test for significance) and is blocked by co-treatment with APC/C inhibitors APCin (40 µM) and proTAME (20 µM) (unpaired *t*-Test or Mann Whitney *U*-test for significance) (**f**). **g**-**i** ‘Non-degradable’ AURKA-Venus is degraded efficiently in response to PROTAC-D: AURKA-Venus-Δ32-66 (ΔA-box) expressed in RPE1 cells (tet-inducible RPE1-FRT/TO pool) is sensitive to PROTAC-D (unpaired *t*-test for significance) (**g**). WT or Ser51D versions of AURKA-Venus transiently electroporated into U2OS cells are equally sensitive to PROTAC-D treatment in either mitotic cells arrested by STLC (**h**) or interphase cells (**i**) (unpaired *t*-test or Mann Whitney *U*-test for significance). Ongoing protein synthesis is likely to mask the degradation rate of Venus-tagged protein in interphase cells.

The effect of PROTAC-D in promoting mitotic exit could potentially be explained by a number of studies showing a role for AURKA in the SAC^27–29^, but could also occur through ‘mitotic slippage’, should there be any non-specific targeting of Cyclin B1 by PROTAC-D in the presence of an active SAC^30^. We tested this possibility using a RPE1-cyclin B1-Venus^KI^ line^31^. Degradation of Cyclin B1-Venus and escape of cells from SAC-induced arrest were both strongly promoted by PROTAC-D (Figure 2d, e), and weakly by Cpd A. However, in contrast to AURKA-Venus^KI^ degradation (Figure 2b), Cyclin B1-Venus^KI^ degradation measured upon PROTAC-D treatment was sensitive to APC/C inhibition (Figure 2f). These results allowed us to conclude that Cyclin B1-Venus degradation in the presence of PROTAC-D is the result of a weakened SAC and that Cyclin B1 is not targeted directly by PROTAC-D.

In further experiments to test that degradation of AURKA in response to PROTAC-D was independent of the well-characterized APC/C-dependent pathway, we used versions of AURKA known to be resistant to APC/C-mediated degradation. AURKA possesses an atypical APC/C degron motif, the so-called A-box, in its N-terminal disordered region^32^. The A-box function appears to be negatively regulated through phosphorylation on Ser51, since phospho-mimic substitution (S51D) at this site blocks mitotic degradation of AURKA^33,34^. We found, using single cell time-lapse degradation assays, that an A-box deleted (Δ32-66) version of AURKA-Venus stably expressed in an RPE-FRT/TO line was efficiently degraded in response to PROTAC-D (Figure 2g). We concluded that PROTAC-mediated degradation of AURKA does not require the substrate motif essential for its canonical degradation, either for ubiquitination, or at any downstream step in substrate processing at the 26S proteasome. We additionally tested the S51D version of AURKA-Venus alongside the WT protein in time-lapse degradation assays, after transient transfection into U2OS cells. We found not only that both WT and ‘non-degradable’ S51D were sensitive to PROTAC-D in mitotic cells (Figure 2h), but that they were also sensitive in interphase cells (Figure 2i), as further confirmation that PROTAC-mediated processing of AURKA for destruction is independent of cell cycle-dependent pathways. We note that the measured rate of degradation is lower in interphase cells than mitotic cells, most likely because degradation is masked by ongoing synthesis (observed as accumulation of the protein in DMSO-treated control cells). We concluded that degradation of AURKA measured in response to PROTAC-D treatment is a direct consequence of PROTAC-D-mediated targeting. Furthermore, since some experiments were carried out using high-level transient expression of transfected constructs (Figure 2h, i), PROTAC-D appears potent enough to clear target protein at significant levels of overexpression in the cell.

### PROTAC-D action is highly selective for AURKA

Next, we asked whether target destruction mediated by PROTAC-D was specific for AURKA. Since the PROTAC target ligand MLN8237 has a degree of selectivity for AURKA over its cellular paralogue AURKB, but is not completely specific (it inhibits AURKB activity at doses of ≥ 50 nM^4^, and the reported selectivity ratio AURKA-TPX2_(1-43)_: AURKB-INCENP_(783-918)_ is approximately 5-fold^6^), we might expect to find some degradation of AURKB in response to a PROTAC carrying the MLN8237 warhead. Furthermore, considering that within the mitotic cell AURKA resides in multi-protein complexes governing its localization and function, we hypothesized that the ‘ectopic’ recruitment of ubiquitination machinery by PROTAC-D might lead to ubiquitination and destruction of AURKA binding partners. Therefore, we examined if PROTAC-D caused a reduction in cellular levels of AURKB, or of two well-known interacting partners of AURKA, TPX2 and TACC3 (Figure 3).

**Figure 3.**
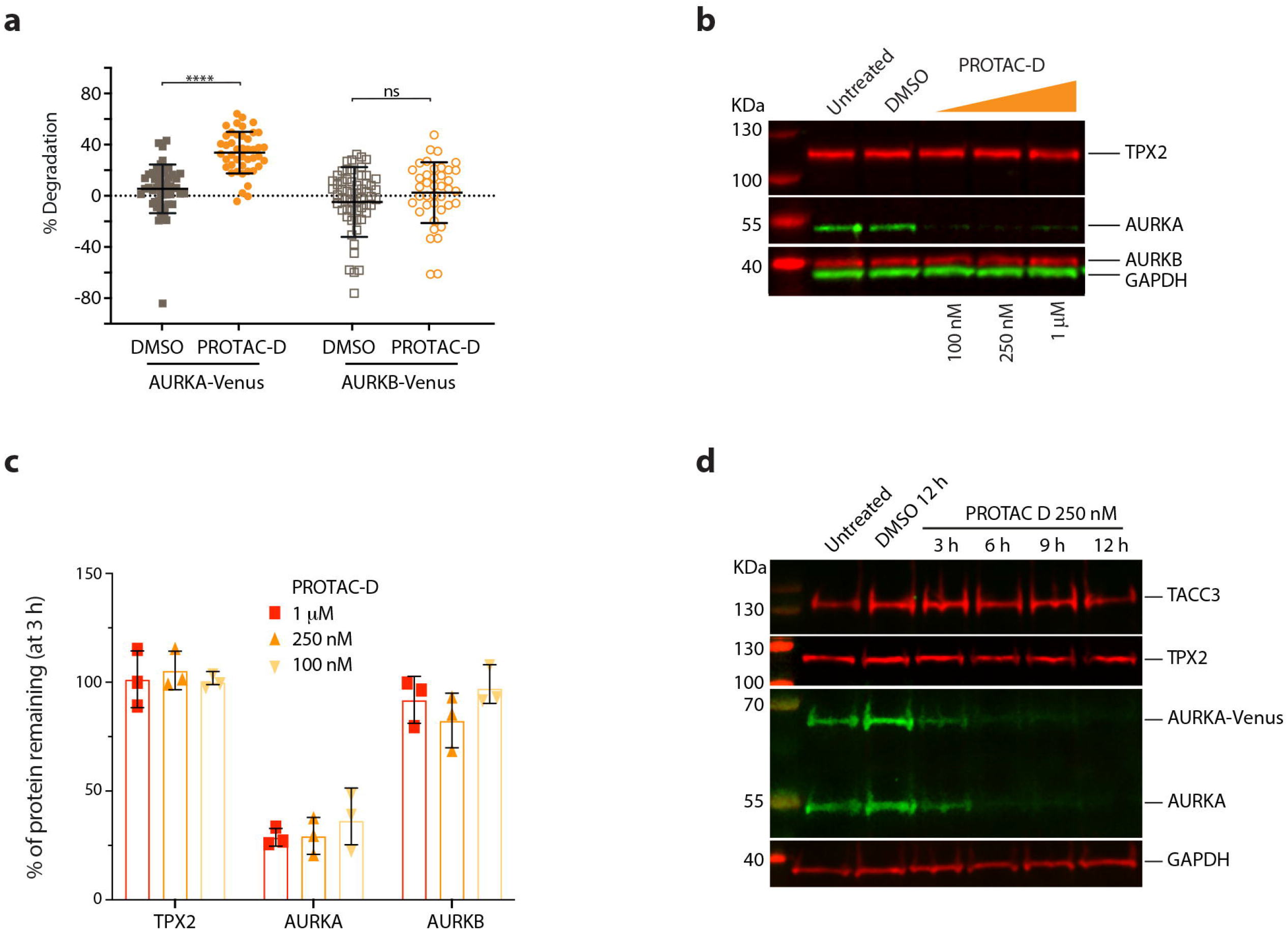
Specificity of AURKA targeting by PROTAC-D. **a** U2OS cells expressing tet-regulated AURKA-Venus and AURKB-Venus were arrested in mitosis with STLC and treated with 250 nM PROTAC-D for 3 hr. Scatter plots with mean values and SDs indicated show percentage degradation of protein in individual cells pooled from three separate experiments, statistical test applied was Mann Whitney *U*-test. **b-d** Endogenous levels of AURKA interactors TPX2 and TACC3, or of AURKB, were examined by quantitative immunoblotting of extracts from cells arrested in mitosis with STLC and treated with PROTAC-D or vehicle control (DMSO). **b** HeLa cells treated for 3 hours, immunoblot representative of data from three identical experiments quantified and presented in **c**, showing mean percentages of protein remaining after PROTAC-D treatment, relative to DMSO treatment, with error bars to indicate SDs. **d** RPE1 AURKA-Venus^KI^ cells treated for 12 hours, immunoblot from one experiment.

Surprisingly, we found that treatment with PROTAC-D caused very little degradation of AURKB-Venus in an inducible U2OS cell line (Figure 3a), or of endogenous AURKB in HeLa cells (Figure 3b, c). We also found no degradation of endogenous TPX2 in mitotically-enriched HeLa cells after 3 hours of treatment with PROTAC-D (Figure 3b, c). TPX2 and TACC3 levels were unchanged in cells treated for up to 12 hr, when endogenous AURKA was no longer detectable (Figure 3d). Therefore PROTAC-D-mediated destruction is highly specific for AURKA. The stability of AURKA binding partners in presence of PROTAC-D suggests that the ubiquitination step is highly specific for the AURKA moiety of mitotic complexes, or alternatively, that only unbound AURKA is targetable by PROTAC-D.

Given the unexpected resistance of AURKB to AURKA PROTAC action, we compared *in vitro* kinase inhibition activities for AURKA and AURKB of PROTACs –D and –DX, Cpd A, and their warhead MLN8237. We found that both of the PROTACs had greater selectivity for AURKA over AURKB than MLN8237 in kinase inhibition assays (fold selectivity of PROTAC-D = 21.6, PROTAC-DX = 23.7, MLN8237 ≥ 8.3) (Table 2), consistent with the lack of AURKB degradation seen in Figures 3a-c. Increased selectivity for AURKA suggests that the increased size and/or complexity of the PROTAC creates new steric parameters influencing target discrimination, and is consistent with published findings from others that the requirement for ternary complex formation in PROTAC action can build a further layer of specificity into drug action^35,36^. Comparing IC_50_ values for inhibition of *in vitro* kinase activity of PROTAC-D and -DX versus MLN8237, we found that inhibition of AURKA kinase activity by the PROTAC molecules is weaker than that mediated by MLN8237 (5-10 fold). Interestingly PROTAC-DX, which has stronger PROTAC activity in comparison to PROTAC-D (Figure 1g), does not have higher activity in this assay (Table 2). This finding is in line with the idea that the efficiency of PROTAC activity is not only impacted by binding affinity to the target or E3 ligase, but also related to efficiency of ternary complex formation between E3 and target protein^37^.

**Table 2:**
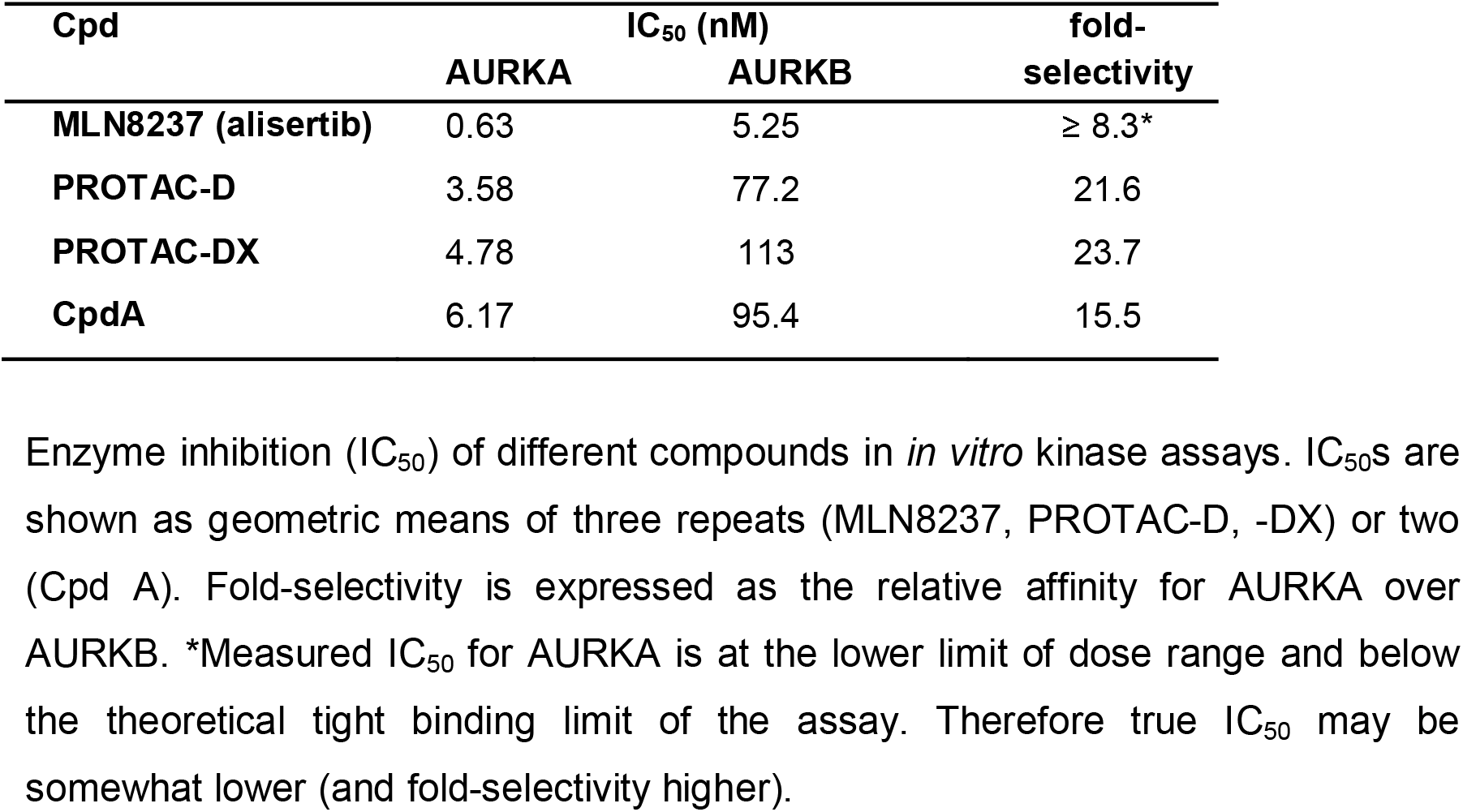
Affinity and selectivity of compounds for AURKA.

### Centrosome-localized AURKA is not sensitive to PROTAC-D

Having validated PROTAC-D as an effective and specific tool for depletion of cellular AURKA, we investigated how PROTAC-mediated AURKA destruction would compare to enzymatic inhibition as a method for down-regulating AURKA functions in mitotic cells. We fixed cell populations synchronized for passage through mitosis and treated for 4 hr with parallel doses of PROTAC-D or MLN8237, or with DMSO as a negative control, and stained them by immunofluorescence (IF) for the presence of AURKA, markers of AURKA activity and tubulin, in order to assess the phenotypic consequences of treatment with different compounds (Figure 4). We looked first at AURKA staining and found that cells treated with PROTAC-D displayed a marked loss of the pool of AURKA associated with the spindle (seen in DMSO-treated controls). However, AURKA was preserved at the centrosomes (Figure 4a). By contrast, treatment with MLN8237 abrogated almost all AURKA localization to centrosomes, consistent with the known role of AURKA activity in centrosome maturation that includes recruitment of AURKA to the pericentriolar material (PCM)^38^. This finding suggested that the centrosome-associated pool of AURKA seen in PROTAC-D-treated mitotic cells would be unexpectedly fully active (that is, neither degraded nor inhibited by PROTAC treatment). We tested this idea by measuring levels of pSer83-LATS2 as a well-known centrosomal marker of AURKA activity, finding that this marker was entirely resistant to PROTAC-D treatment (at doses sufficient to deplete most of the cellular pool of AURKA), whilst responding in dose-dependent fashion to MLN8237 (Figure 4b,c).

**Figure 4.**
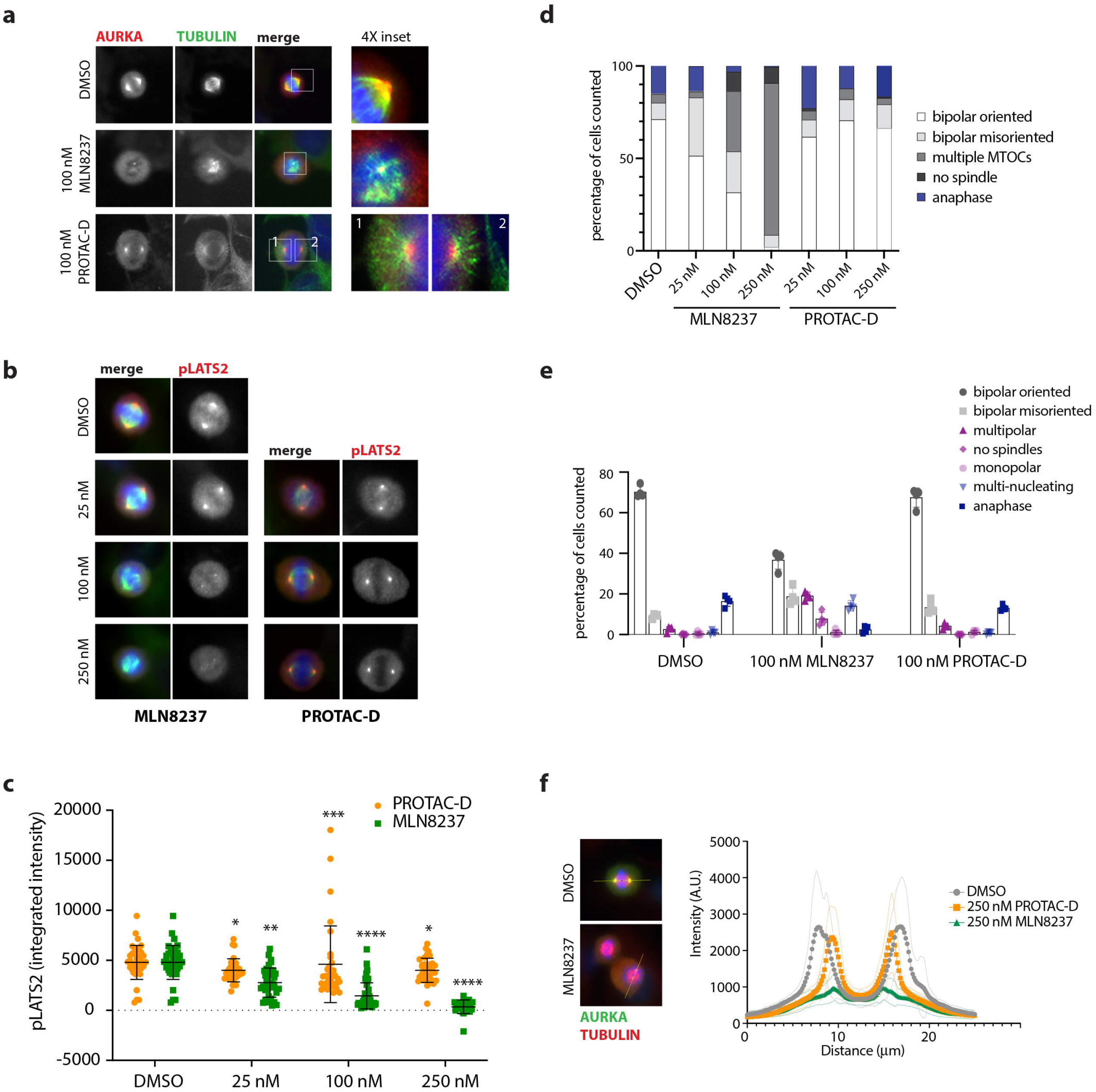
AURKA remains active at centrosomes upon PROTAC treatment. **a-c** Immunofluorescence analyses of U2OS cells synchronized through mitosis and treated for 3 hours with MLN8237, PROTAC-D or DMSO vehicle control. MLN8237 is inhibitory to MT nucleation throughout the mitotic cell. AURKA staining is retained at the centrosomes after PROTAC-D treatment (**a**) whilst a marker for active AURKA at centrosomes, p(Ser83)LATS2, persists after PROTAC-D treatment but not MLN8237 treatment (**b**,**c**). **c** Measurement of centrosomal pLATS2 staining from immunofluorescence images after cytoplasmic background subtraction (Kruskal-Wallis multiple ANOVA, and Dunn’s post-hoc multiple comparison test to DMSO for significance). **d**,**e** Mitotic phenotypes scored from TPX2/DAPI staining of fixed cells (categories illustrated in Supplementary Figure S3). **d** Dose-dependence of phenotypes using pooled data from 3 coverslips for each condition from a single experiment. **e** Mitotic categories scored at a single dose (100 nM), using data from 2 coverslips each from 3 independent repeats of the experiment, mean values ± SDs for each experiment are plotted. **f** Line-scans through fixed mitotic cells stained with pLATS2 (as shown in Figure 4B), to show both poles of the spindle (visible as maxima of staining intensity), reveal reduced pole-pole distance after PROTAC-D treatment. Traces show mean values from n ≥ 22 individual mitotic spindles, with fainter traces above and below representing SDs.

Given the >5-fold difference in enzyme inhibition of PROTAC-D and MLN8237 (Table 2) and the likelihood that the intracellular dose of PROTAC-D is limited by its size and solubility, we examined the phenotypic consequences of treatment over a 10-fold range of doses of both compounds, scoring mitotic figures according to the categories illustrated in Supplementary Figure S3. Dose-response to MLN8237 treatment is characterized by progression from spindle orientation defects at low doses to spindle assembly defects (multipolar spindles, ‘small’ spindles) at intermediate doses, to lack of MT nucleation at a dose of 250 nM (Figure 4d and as previously described^4^). We were surprised to find that PROTAC-D-treated cells showed none of these defects (Figure 4d, e). Even at the highest dose tested (250 nM), we did not see the orientation defects characteristic of low dose inhibition of AURKA activity^4,39^. Instead, we observed that the mitotic spindles were shorter in length after PROTAC treatment. Distribution of the centrosomal pLATS2 staining shown in Figure 4b confirms that the pole-to-pole distance of correctly oriented bipolar spindles is reduced (Figure 4f).

### PROTAC-D and MLN8237 mediate distinct effects in mitotic cells

Our finding of ‘short spindles’ was reminiscent of the previously reported finding that specific perturbation of AURKA binding to TPX2 controls spindle length independently of any effect on assembly^11^ which can occur under the influence of the centrosomal AURKA pool. Therefore, we hypothesized that PROTAC-D had selectively depleted the TPX2-associated pool of AURKA to eliminate the chromosome-centred MT nucleation pathway whilst leaving the centrosomal pathway untouched. We decided to test this idea using a modified cell synchronisation assay that would better allow us to compare the roles of kinase inhibition and target degradation in mitotic cells independent of their different effects on AURKA-dependent centrosome maturation. We pre-synchronised cells at metaphase by release of cells arrested for 24 hours in Thymidine into APCin/proTAME for 6 hours. We then treated metaphase-arrested cells with different doses of MLN8237, PROTAC-D and Cpd A for 3 hours before fixing them for IF analysis. We first examined the pT288 epitope as a direct readout for inhibition of AURKA, finding as expected that PROTAC-D and Cpd A show weaker inhibition of AURKA activity than MLN8237. Importantly, the extent of inhibition by PROTAC-D and Cpd A is similar, confirming that both are able to access the centrosomal pool of AURKA (Figure 5a,b) and enabling us to reason that comparing the effects of these compounds would reveal phenotypes arising from degradation of AURKA, distinct from those arising purely out of kinase inhibition.

**Figure 5.**
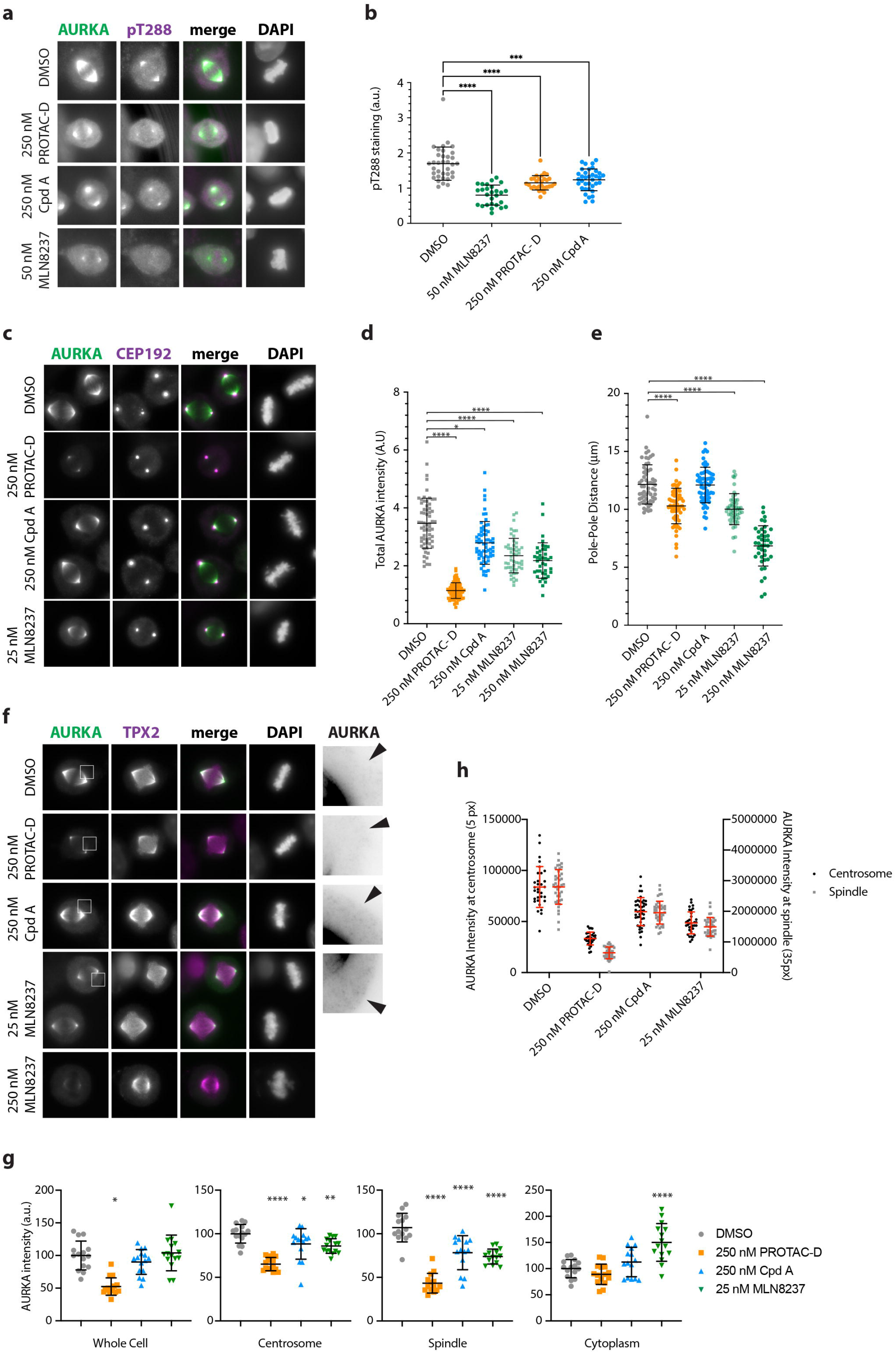
PROTAC specifically depletes AURKA on the mitotic spindle. U2OS cells were released from single thymidine block into APCin and ProTAME to arrest at metaphase and pre-treated for 3 hours with DMSO, PROTAC-D, Cpd A or MLN8237 before fixation. Cells were then stained for AURKA, pT288-AURKA and DAPI **(a, b)**, AURKA, CEP192, and DAPI (**C-E**) or AURKA, TPX2 and DAPI **(f-h)**. All data shown are representative of two identical experiments. **a** Examples of cells imaged under different drug treatments and used for measurement of pT288-AURKA intensity at spindle poles. **b** Data are shown as scatter plots with means and SDs indicated and were tested for statistical significance by ordinary one-way ANOVA with Dunnett’s post-hoc test for significance to DMSO. **c** Examples of cells imaged under different drug treatments and used for measurement of total AURKA intensity after background subtraction (**d**) or measurement of spindle pole separation as distance from one CEP192-marked centrosome to the other (**e**), shown as scatter plots with means ± SDs indicated and tested by Kruskal-Wallis multiple ANOVA, and the Dunn’s post-hoc multiple comparison test to DMSO. **f** Examples of cells imaged under different drug treatments and used to compare AURKA levels on the spindle and at the spindle poles. Insets indicated with white boxes in AURKA channel are enlarged 4X in righthand panels and have contrast increased to reveal cytoplasmic levels of AURKA (arrowheads). **g**,**h** Integrated AURKA intensities were measured within circles of fixed diameter centred on the centrosomes, spindle or cytoplasm to show the effect of PROTAC-D treatment on different pools of AURKA and reveal that loss of AURKA from spindle and centrosomes in control drug treatments is reflected in increased cytoplasmic levels (**g**) and that PROTAC-D treatment causes greater loss of AURKA from the spindle than from the centrosomes (**h**). Scatter plots with means ± SDs indicated show measurements from individual cells from a single experiment. For plot **g**, values were normalized to mean DMSO levels and tested by ordinary one-way ANOVA with Dunnett’s post-hoc test for significance to DMSO. For plot **h**, raw values were plotted using separate Y axes for centrosome and spindle values (integrated over circles of 5 and 35 pixel diameters, respectively).

Next, we stained cells for AURKA and its interactors CEP192 and TPX2 (Figure 5c-h). Similar levels of CEP192 at centrosomes after the different treatments confirmed that centrosome maturation had occurred in a large fraction of the cellular pool of metaphase cells (Figure 5c). Quantification of AURKA levels in these cells showed the total cellular pool of AURKA reduced more than three-fold after PROTAC-D treatment (Figure 5d). Measured AURKA levels were also somewhat lower (by about 30%) after treatment with Cpd A or with low (25 nM) or high (250 nM) doses of MLN8237. Since we have found that these treatments do not affect endogenous AURKA levels, nor AURKA-Venus levels in intact cells, we assumed that the reduced AURKA levels seen in IF reflected loss of AURKA in the fixation step, that could be a consequence of reduced interaction with the mitotic spindle. Indeed, MLN8237 and TPX2 may compete with each other for AURKA binding^40^ (see Discussion). We measured pole-pole distances in this experiment and found them reduced by PROTAC-D treatment. AURKA inhibition with 25 nM MLN8237 also gave rise to short spindles, whereas Cpd A had no effect on spindle length (Figure 5e). We concluded that Cpd A and PROTAC-D both bind too weakly to AURKA to significantly inhibit its activity, and that the short spindle phenotype seen after PROTAC-D treatment depends on destruction of AURKA by PROTAC-D. Consistent with this conclusion, we observed that PROTAC-D alone of the three treatments removed both cytoplasmic and spindle pools of AURKA (Figure 5f). Kinase inhibition mediated by 25 nM MLN8237 or 250 nM Cpd A caused some loss of signal from the spindle, but also increased cytoplasmic levels of AURKA (Figure 5f, g). Comparison of AURKA pixel intensities in fixed areas on the centrosome or neighbouring spindle confirmed that depletion of the spindle signal was greater than at the centrosome (Figure 5h). We concluded from our data that PROTAC-D preferentially depletes the pool of AURKA that associates with TPX2 to govern mitotic spindle length. Moreover, because kinase inhibition assays indicate that Cpd A and PROTAC-D bind AURKA with equivalent affinity (Figure 5b, Table 2), we concluded that the short spindle phenotype seen after PROTAC-D - but not Cpd A - treatment (Figure 5e) depends on destruction of AURKA protein.

### Selective depletion of distinct target pools by PROTAC-D

We investigated further why PROTAC-D treatment led to selective depletion of the spindle-associated pool of AURKA, hypothesizing that this could be a consequence of conformation-dependent targeting by the PROTAC, with the preferred target being either the TPX2-bound pool or a free pool of AURKA (provided this turns over faster with the TPX2-bound pool than the centrosomal pool). We tested this idea by using conformational mutants of AURKA-Venus. AURKA-S155R is a version of AURKA showing reduced interaction with TPX2 (and therefore weak localization to the mitotic spindle), whilst its centrosomal localization is maintained^41^. The non-catalytic N-terminal IDR of AURKA is known to mediate interaction with a number of binding partners^42–44^, so N-terminally truncated versions of AURKA, 68-403 (Δ67) and 128-403 (Δ127), are predicted to show reduced interactions with binding partners, including a potential auto-inhibitory interaction with the kinase domain^23,45^.

We found that in RO3306-arrested interphase cells, overexpressed S155R and wild-type versions of AURKA-Venus showed a similar pattern of targeting by PROTAC-D. However in cells arrested in prometaphase with STLC, where TPX2 is a major binding partner of AURKA, S155R was more rapidly degraded than the wild-type, consistent with the idea that PROTAC-D preferentially depletes free AURKA in mitotic cells (Figure 6a). The N-terminally truncated version of AURKA Δ127 was insensitive to PROTAC-D, and in the absence of PROTAC treatment accumulated strongly (Supplementary Figure S4) and we concluded that it probably lacks substrate lysines and/or IDR required for ubiquitination or processing at the 26S proteasome. The Δ67 version of AURKA-Venus was, however, more sensitive to PROTAC-D than the full-length protein under all conditions tested (Figure 6b, Supplementary Figure S4). This finding is consistent with the idea that interactions mediated through the N-terminal IDR impede PROTAC-D-mediated degradation of the target.

**Figure 6.**
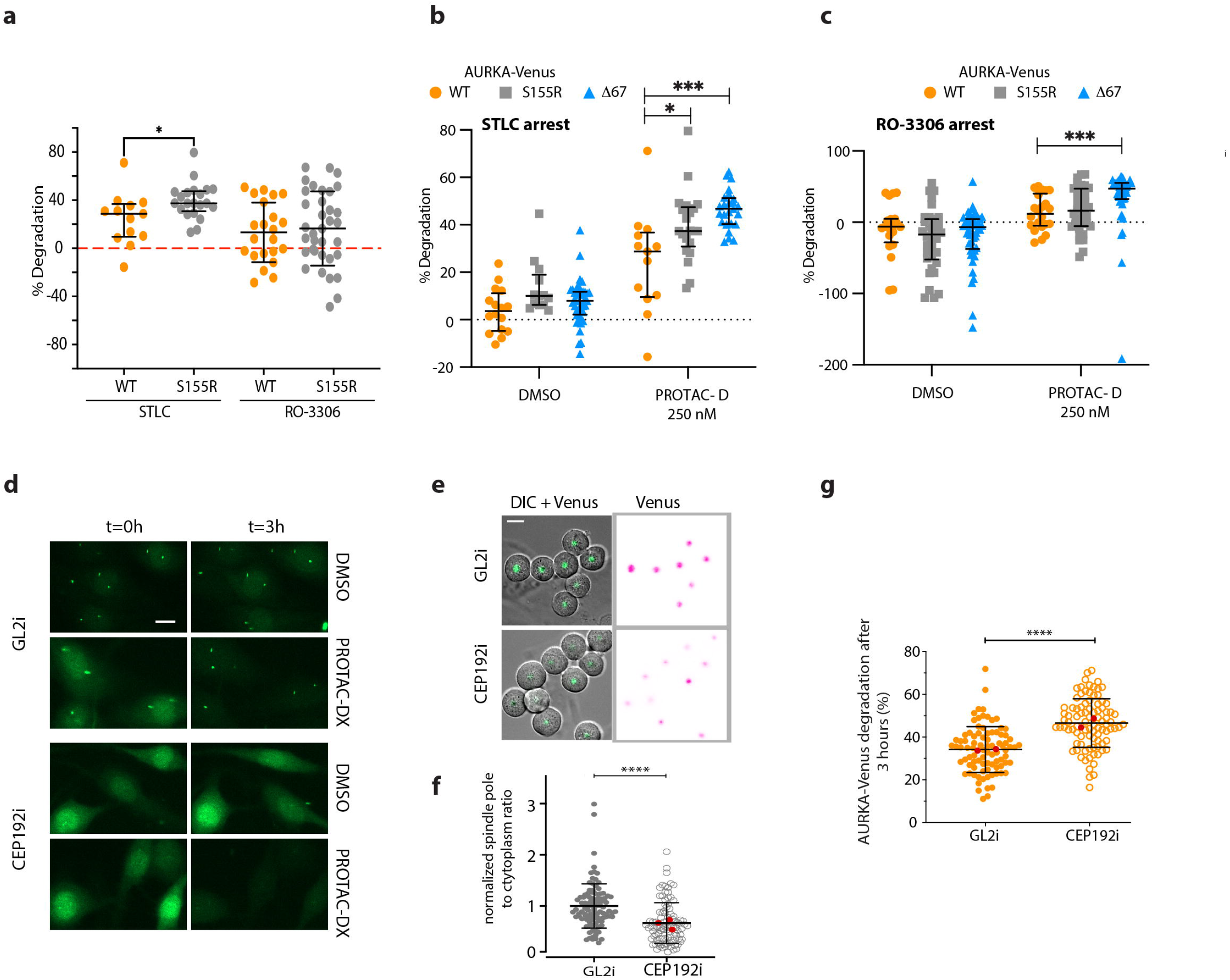
Differential targeting of AURKA pools. **a-c** Differential degradation of AURKA-Venus variants in response to PROTAC-D. U2OS cells were transiently transfected with different versions of AURKA-Venus and treated for 16 hours with RO-3306 to arrest them in G2 phase, or with STLC to arrest them in mitosis. Cells were then treated with 250nM PROTAC-D or DMSO as vehicle control and subjected to fluorescence timelapse microscopy over 3 hours. Venus levels in whole cells were measured at the start and end of each experiment to calculate percentage protein remaining for each cell. Scatter plots show data from individual cells to compare wild-type and S155R versions (**a**) or wild-type, S155R and Δ67 versions under conditions of mitotic (**b**) and interphase (**c**) arrest. Bars and whiskers represent mean values and SDs and are pooled from three repeats of the same experiment (unpaired *t*-test, n ≥ 10). **d-g** RPE1-AURKA-Venus^KI^ cells were transfected for 48 hours with siRNA against CEP192 (CEP192i) or control siRNA (GL2i) then either arrested in G2 phase with RO3306 (**d**) or arrested in mitosis with STLC (**e-g**) for 16 hours before treatment with PROTAC-DX for 3 hours. **D** Panels from start and end of experiment show AURKA-Venus preserved at centrosomes in control cells even following PROTAC treatment, but absent from centrosomes in CEP192i (See Supplementary Figure S5 for quantification of this experiment). **e** Panels from start of experiment in STLC-arrested cells, showing reduced localization of AURKA to the single spindle pole in each cell following CEP192i, as quantified in **f**: Mean pixel intensities of AURKA-Venus were measured within a region of interest of fixed diameter around poles and in the cytoplasm, and are expressed as the ratio of spindle pole:cytoplasm after normalization of each dataset with the mean value of the control sample. Data from individual cells in 3 independent repeats of the experiment were pooled, with the mean value of each repeat plotted in red and tested for statistical significance using Mann Whitney *U*-test. **g** In mitotic-arrested cells AURKA-Venus^KI^ showed strongly enhanced degradation after CEP192 knockdown in live cell degradation assays: Plots show percentage degradation in individual cells after 3 hours treatment and contain pooled data from 2 independent experiments with mean values from each experiment plotted in red (Mann Whitney *U*-test). A third independent experiment was carried out, showing the same result but with response to PROTAC reduced in all conditions, with the pooled data shown in Supplementary Figure S6.

In order to confirm that subcellular pools of wild-type AURKA also showed differential sensitivity to PROTAC-D, we experimented instead on RPE1-AURKA-Venus^KI^ cells treated with siRNA against CEP192 (CEP192i) to displace AURKA from the centrosomes^10^. Displacement of AURKA-Venus was readily observed in cells arrested in G2 phase with RO3306, when AURKA-Venus^KI^ expression is high (Figure 6d, quantified in Supplementary Figure S5). We tested AURKA-Venus degradation in response to PROTACs under these conditions and measured slightly increased degradation of AURKA-Venus in single cells or by immunoblot (Supplementary Figure S5). We then tested the effect of CEP192i on mitotic cells, when a much larger pool of AURKA is normally recruited to centrosomes. In STLC-arrested cells we found that AURKA-Venus^KI^ delocalized from centrosomes after CEP192i (Figure 6e, f) was more readily degraded in response to PROTAC-D (Figure 6g). We concluded that the centrosome localized pool of AURKA-Venus shows reduced sensitivity to PROTAC-D.

### Inhibition of cytoplasmic AURKA activity by PROTAC-D

Recent studies of AURKA have highlighted that interphase functions may contribute to its tumorigenic activity. One interphase function is regulation of the mitochondrial network and our recent work has shown that excess AURKA present in interphase FZR1^KO^ cells causes fragmentation of the mitochondrial network^24^. This activity of AURKA, promoted through stabilization of the protein, represents an interesting context for testing PROTAC strategies, so we tested the action of PROTAC-DX on mitochondrial fragmentation in FZR1^KO^ cells. We treated FZR1^KO^ cells with PROTAC-DX and Cpd A and found that PROTAC-DX, but not Cpd A, restores mitochondrial morphology (Figure 7a,b). Therefore PROTAC-DX is able to prevent interphase activity of AURKA in a manner that depends on destruction of the protein. We conclude that PROTAC-mediated clearance is an efficient strategy for downregulating excess AURKA activity in interphase cells.

**Figure 7.**
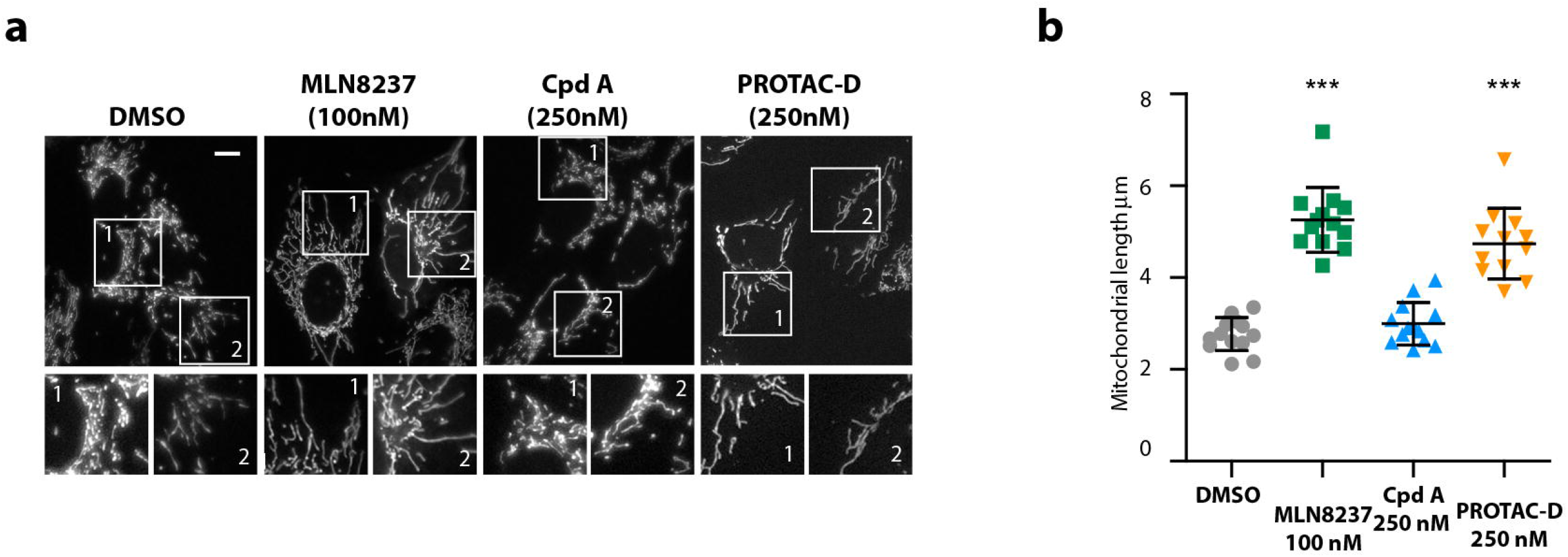
PROTAC-D treatment restores the mitochondrial network in FZR1^KO^ cells. U2OS FZR1^KO^ cells, in which mitochondria are highly fragmented due to the presence of excess AURKA^24^, were treated for 3 hours with DMSO or with MLN8237, Cpd A or PROTAC-D at the doses indicated. Cells were then stained with MitoTracker Red™, imaged and analysed for mitochondrial fragment lengths as described in materials and methods. **a** Representative images of cells analysed. Scalebar, 10μm. **b** Scatter plots showing mean mitochondrial lengths, mean and SDs under different drug treatments. Each datapoint represents the mean value from 30 mitochondria per cell, n ≥ 12 cells, with results tested for significance using Mann Whitney *U*-test. The data are from a single experiment that is representative of two identical repeats of the same experiment.

## Discussion

We have described a small molecule that acts as a specific degrader of AURKA to clear endogenous, exogenous GFP-tagged, or overexpressed protein from the cell. Amongst the molecules we tested, successful degraders were CRBN-specific. Although we have not formally excluded the possibilities that VHL is insufficiently active in U2OS cells to generate degradation-competent ubiquitin conjugates in response to Cpd A, or that the four linker constructs we tested all occluded ternary complex formation between AURKA and VHL, our observation that the same four linker constructs were all able to support PROTAC activity directed to CRBN are in line with the published finding that protein-protein interaction surfaces of CRBN are more favourable to stable ternary complex formation than the equivalent surfaces of VHL^37^. As widely reported in the literature, PROTACs have poor solubility and cell permeability. Therefore we also considered the possibility that our VHL-directed compounds are less cell permeable than their CRBN-directed matching compounds, by testing our VHL-directed ‘control’ compound, Cpd A, for its ability to inhibit AURKA activity both in cells as well as *in vitro*. We found that PROTAC-D and Cpd A showed similar activity in both assays (Figure 5a, Table 2).

Compounds showing PROTAC activity against AURKA were several-fold less potent than their MLN8237 warhead in inhibiting AURKA activity *in vitro*, consistent with reduced affinity for their target. Indeed, we found that the partial inhibition of AURKA activity in response to PROTAC-D treatment (pT288-AURKA staining, Figure 5) was insufficient to reduce AURKA function at the centrosomes (pS83-LATS2 staining, Figure 4), and that non-centrosomal AURKA was insensitive to Cpd A inhibition (since mitotic spindle assembly was unaffected). Therefore we concluded that binding of PROTAC-D is weak enough, and/or the molecule present at sufficiently low intracellular levels, to achieve targeted degradation of AURKA in absence of functional consequences to the inhibition of AURKA kinase activity, and without exhibiting the hook effect characteristic of heterobifunctional ligands^46^. The question of the intracellular effective dosage of PROTACs remains an elusive parameter.

We found that the activity of CRBN-directed molecules correlated with linker length but was independent of the affinity of the compound for its AURKA target. Therefore, it is likely that our longer linkers promote the assembly of productive ternary complexes by bringing together AURKA and CRBN in an orientation that allows the E3 complex to ubiquitinate AURKA at appropriate lysine residues. The physical properties of the linker are critical parameters in PROTAC activity, and further optimisation of PROTAC-DX could include different linker patterns to alter linker flexibility, as well as lengths.

We observed that clearance of wild-type AURKA from the cell is less efficient than that mediated by its cognate E3, APC/C-FZR1 (t_1/2_ ∼ 100 min vs t_1/2_ ∼ 45 min^18^). We speculate that even with further optimization, it seems unlikely that any PROTAC would eliminate AURKA faster than its cognate pathway, since the rate-limiting step for degradation of many ubiquitinated substrates is not recruitment to the 26S proteasome, but determinants of processing that are partly substrate-specific (such as unfolding of substrate at the proteasome) and partly determined by the configuration of ubiquitin chains^47^. Indeed, a recent study found that the presence of unstructured regions determines the PROTAC mediated degradation of VHL-directed substrates^48^. However, we found that PROTAC-D could clear versions of AURKA resistant to APC/C-mediated degradation (through mutation or removal of the essential N-terminal degron) as efficiently or better than the wild-type AURKA. We conclude from this that the position and topology of ubiquitin chains assembled on AURKA by CRBN and APC/C-FZR1 are likely to be very different, and that PROTACs are promising tools for targeting mutant versions of cellular proteins that have acquired resistance to canonical degradation pathways.

We also observed that different cellular pools of AURKA substrate were differentially targeted by PROTAC treatment, since in mitotic cells the spindle-associated fraction of AURKA was eliminated whilst the centrosome fraction was preserved (Figure 8). Since the centrosomal pool of AURKA retained its activity, spindle assembly was buffered against the loss of the chromatin-associated TPX2-activated AURKA pool and the observed phenotype of PROTAC-D treatment in mitotic cells is therefore shortened spindles, consistent with a previous study of cells engineered to express a non-AURKA-binding version of TPX2^11^. Similarly, in interphase cells we observed that PROTAC-D treatment efficiently cleared the non-centrosomal pool of AURKA, but that centrosomal AURKA was preserved. Delocalization of the centrosomal pool through siRNA-mediated depletion of CEP192 promoted clearance of the total cellular pool of AURKA-Venus by PROTAC-D. Since centrosomal AURKA is efficiently inhibited by MLN8237, we would expect it to be accessible to bind MLN8237-derived PROTAC molecules. One explanation for its inaccessibility to PROTAC-D action could be that centrosomal localization, or the interaction with CEP192, block recruitment of CRBN or another component of the E3 complex required for ubiquitination of its target. Alternatively, there may be deubiquitinase enzymes active at the centrosomes that act to stabilize ubiquitinated AURKA.

**Figure 8.**
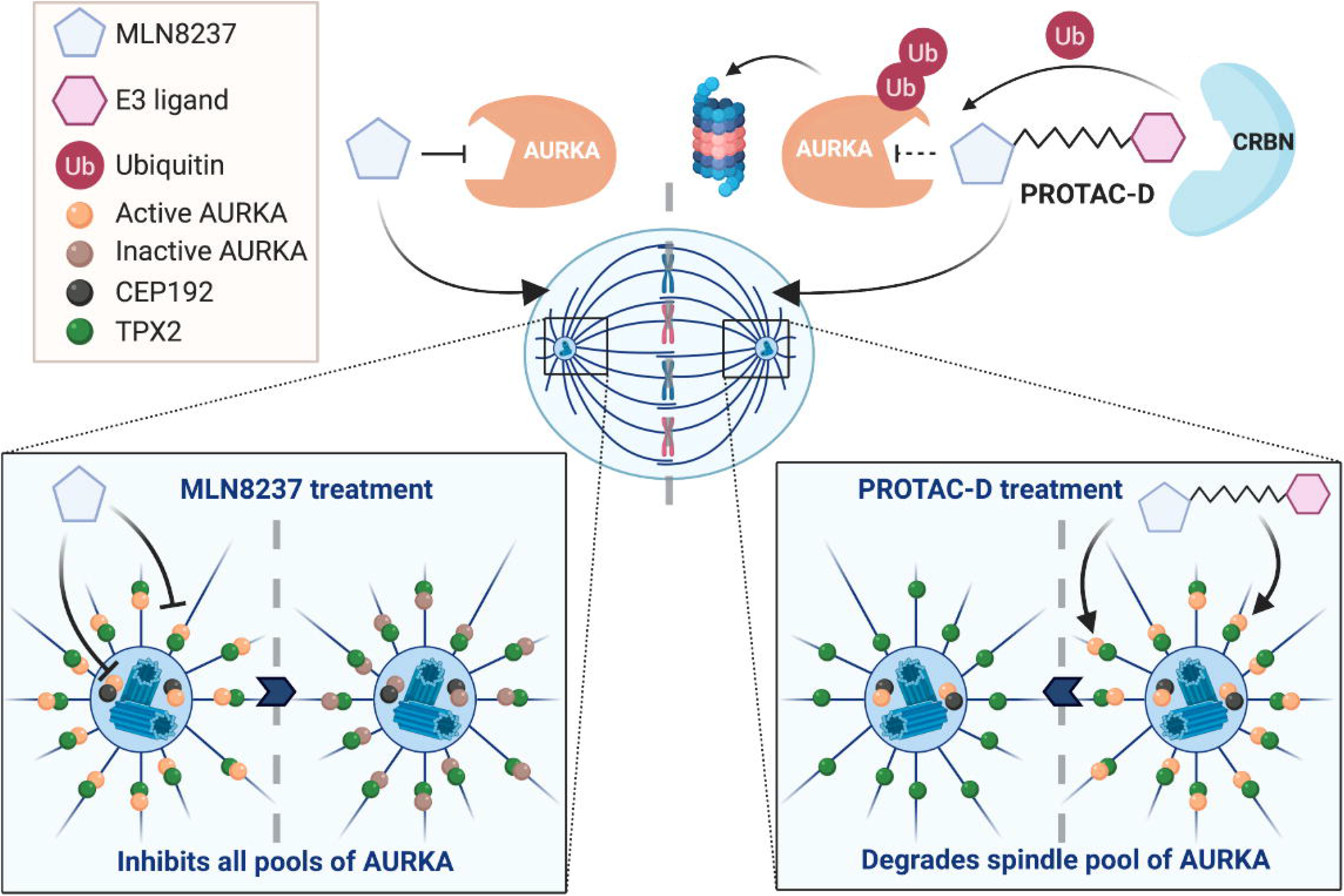
Targeting of non-centrosomal AURKA functions by PROTAC-D. Schematic to illustrate the distinct actions of MLN8237 and PROTAC-D on AURKA pools in mitotic cells. Both centrosomal and spindle pools of AURKA are inhibitable by MLN8237, whereas PROTAC-D clears the spindle pool of AURKA whilst leaving active AURKA at the centrosome. Figure made in ^©^BioRender (biorender.com).

Our results point to differential accessibility of subcellular pools of substrate, governed by substrate conformation, binding partners, or localization in compartments more or less accessible to PROTAC action, a phenomenon that has not previously been described for PROTAC agents acting via CRBN. Our finding of localized response to PROTAC-D is in contrast with treatment by the AURKA inhibitor alisertib, which promotes a clear dose-dependent depletion of pLATS2, a marker of AURKA activity at the centrosomes.

Given the complex conformational and spatial regulation of AURKA^14,16^ we tested for conformation-specific targeting of the kinase using different versions of AURKA-Venus. The conformational dynamics of AURKA are strongly constrained through interaction with TPX2^40,49,50^, which favours the so-called ‘DFG-In’ active confirmation. A recent study from the Levinson group suggests that different functional pools of the kinase possess distinct conformational properties that modulate interactions with inhibitors, with alisertib/MLN8237 shown to be a ‘Type 1’ inhibitor that promotes the inactive DFG-Out state, and TPX2 shown to oppose DFG-Out inducers, exhibiting negative cooperativity on binding with MLN8237. In the DFG-Out state, the active site is more open (i.e. more accessible to MLN8237 or PROTAC-D) and in this model, a version of AURKA impaired in TPX2 binding (S155R) should be more strongly degraded in response to PROTAC-D in presence of TPX2. Indeed, we found some difference in mitotic, but not interphase cells, consistent with a competition model whereby AURKA bound by TPX2 is less accessible to PROTAC-D binding. However, it is also likely that AURKA in complex with binding partners is less accessible for ternary complex formation with CRBN and other components required for efficient ubiquitination in response to PROTAC-D. We tested this idea with truncation of the non-catalytic domain of AURKA to reduce interaction with some binding partners whilst not affecting the ability of PROTAC-D to access the catalytic domain. Δ67 was degraded more efficiently than the full-length protein under all conditions tested, consistent with the idea that binding partners influence the ability of PROTAC-D to bring about degradation of its target.

AURKA is of strong interest as a therapeutic target for various cancers, but despite extensive testing in clinical trials, alisertib has yet to reach the clinic. Our study is the first to describe a degrader compound that shows specificity for different subcellular pools of AURKA, raising the possibility of developing PROTACs to fine-tune the activity of AURKA (and other targets that have shown disappointing clinical results) to produce cellular phenotypes that are potentially more desirable in pharmacological or therapeutic contexts. For example, alisertib-derived PROTACs could be used to target cytoplasmic functions of AURKA without inducing mitotic errors that are consequence of inhibiting AURKA function at the centrosome.

## Materials and Methods

### Cell culture and synchronization

U2OS and HeLa cells were cultured in DMEM (Thermo Fisher Scientific) and hTERT-RPE-1 cells in DMEM:F12 mix, supplemented with 10% FBS, 200 µM Glutamax-1, 100 U/ml penicillin, 100 µg/ml streptomycin, and 250 ng/ml fungizone. All cells were grown in humidified atmosphere at 37°C with 5% CO_2_. RPE-1 AURKA^KI^ cells, RPE-1 AURKA^TO^ cells, Tet-Off cell lines U2OS-AURKA-Venus and U2OS-AURKB-Venus, have all been previously described^25,52^.

For assaying live cell degradation of AURKA-Venus^KI^ and AURKA-Venus^TO^ in mitotic arrested cells, 1.5 × 10^4^ RPE1 AURKA-Venus^KI^ cells were seeded per well in 8-well slides (Ibidi GmbH) and treated for 16 hr with 10 µM S-trityl L-cysteine (STLC) (Tocris Bioscience) prior to PROTAC treatment.

For assaying live cell degradation of AURKA-Venus^TO^ in G_2_ arrested cells, 1.5 × 10^4^ RPE1 AURKA^TO^ cells were seeded per well in 8-well slides (Ibidi GmbH) and treated for 16 hr with 10 µM RO3306 (Tocris Bioscience) prior to PROTAC treatment.

For assaying degradation by immunoblot of cell extracts, 2 × 10^5^ AURKA-Venus^KI^ cells were seeded in 6-well plates prior to 16 hr STLC treatment and addition of test compounds.

Cells for immunofluorescence were seeded on glass coverslips and enriched for the population of mitotic cells by release from a single 24 hr block with 2.5 mM Thymidine. Cells were fixed 10 hr after release, to include the time of treatment with test compounds.

For assaying mitochondrial fragmentation, U2OS FZR1^KO^ cells^24^ seeded on 8-well Ibidi slides were incubated for 15 minutes at 37°C in MitoTracker Red^®^ CMXRos(Thermo Fisher Scientific) as per manufacturer’s instructions.

### Drug treatments

AURKA PROTACs and pomalidomide were synthesised in-house and used at ≤ 1 μM. Detailed chemical synthesis of Compounds A-H is described in Supplementary Methods. Aurora A kinase inhibitor MLN8237 (Stratech, Ely, UK) was used at ≤ 1μM, MG132 (Alfa Aesar) at 42 µM, RO3306 (Tocris Bioscience) at 10 µM, APCin (Bio Techne) at 20 µM and ProTame (R&D Systems) at 40 µM.

### Cell transfection

Cells were transfected with 1 µg of plasmids using electroporation with Neon Transfection System (Thermo Fisher Scientific) using the following parameters: pulse voltage 150 V, pulse width 10 ms, and 2 pulses total on the transfection device according to the manufacturer’s protocol. AURKA and AURKB plasmids used were expressed with C-terminal Venus tags in pVenus-N1 vector. Δ32-66, S51D, S155R Δ67 and Δ127 versions of AURKA were generated by PCR mutagenesis, with cloning maps available on request. CEP192 knockdown was achieved by transfecting the oligo duplex: 5’-GGAAGACAUUUUCAUCUCUtt-3’ and 5’-AGAGAUGAAAAUGUCUUCCtt-3’ (Sigma).

### Immunoblotting

Cell extracts were prepared in NuPage (Invitrogen) SDS sample buffer with 100 µM DTT, Extracts were syringed and boiled prior to electrophoresis on NuPage precast 4-12% Bis-Tris SDS-PAGE gels (90 min, 150 V, 80 W). Proteins were transferred on to Immobilon-FL PVDF (Sigma) membrane using a wet transfer XCell IITM Blot Module system (120 mins, 30 V, 80 W). Blocking and incubations were performed in phosphate-buffered saline (PBS), 0.1% Tween-20, 5% low-fat milk (TBST and 3% BSA for phosphoantibodies) either overnight at 4 °C or for 1 hour at room temperature. Signals were quantified by enhanced chemiluminescence detection, or using fluorophore-conjugated secondary antibodies, scanned on an Odyssey® Imaging System (LI-COR Biosciences). The uncropped raw tif files for each blot are assembled in Supplementary Figure S7.

Primary antibodies for immunoblot were as follows: AURKA mouse mAb (1:1000; Clone 4/IAK1, BD Transduction Laboratories), phospho-Aurora A (Thr288)/Aurora B (Thr232)/Aurora C (1:1000; clone D13A11 XP® Rabbit mAb, Cell Signalling), rabbit polyclonal TPX2 antibody (1:1000; Novus Biological), AURKB rabbit polyclonal antibody (1:1000; Abcam ab2254), mouse mAb Cyclin B1 (1:1000; BD 554177), rabbit polyclonal beta-tubulin (1:2000; Abcam ab6046), GAPDH rabbit mAb (1:400; Cell Signaling Technology #2118), TACC3 rabbit polyclonal antibody (1:1000; gift from F. Gergely), CEP192 affinity-purified rabbit polyclonal antibody (1:1000; Gift from L. Pelletier^53^).

Secondary antibodies used were Polyclonal Goat Anti-Rabbit or Polyclonal Rabbit Anti-Mouse (1:1000) HRP-conjugated (Dako Agilent), or IRDye® 680RD (1:20,000)- or 800CW (1:10,000)-conjugated for quantitative fluorescence measurements on an Odyssey® Fc Dual-Mode Imaging System (LICOR Biosciences). IRDye® conjugated antibodies were prepared in PBS, 0.1% Tween-20, 5% FBS, 0.01% SDS.

### Immunofluorescence

Cells were seeded at 2 × 10^4^ onto glass coverslips and then fixed with cold 100% methanol (−20°C), permeabilized and blocked with 3% bovine serum albumin (BSA) and 0.1% Triton X-100 in PBS (blocking buffer) for 15 min at room temperature. Cells were washed 3 times in PBS with 0.1% Triton X-100 for 5 min each prior to 1 hour incubation with primary antibodies diluted in blocking buffer at room temperature in a humidity chamber. Slides were then washed 3 times again in PBS with 0.1% Triton X-100 for 5 min each before incubation with secondary antibodies diluted in blocking buffer for 45 min at room temperature in a humidity chamber. DNA was stained with Hoechst-33342 (1 µg/mL) and coverslips were mounted with Prolong Gold antifade reagent.

Primary antibodies used for immunofluorescence were as follows: AURKA mouse mAb. (1:1000; Clone 4/IAK1, BD Transduction Laboratories), AURKA rabbit polyclonal (1:1000; Abcam ab1287), PLATS2 mouse mAb (1:1000; Clone. ST-3B11, Caltag Medsystems), TACC3 rabbit polyclonal antibody (1:1000; gift from F. Gergely), TPX2 rabbit polyclonal (1:1000; Novus Biological)

CEP192 affinity-purified rabbit polyclonal antibody (1:1000; Gift from L. Pelletier), beta-tubulin rabbit polyclonal (1:1000; Abcam ab6046), beta-tubulin mouse mAb (1:300; Sigma T4026)

Secondary antibodies used were: Alexa Fluor 488 anti-mouse and Alexa Fluor 568 anti-rabbit (Thermo Fisher Scientific).

### Microscopy

All images were acquired on automated epifluorescence imaging platforms based on Olympus IX81 or IX83 inverted microscopes (Olympus Life Science, Southend-on-Sea, UK) with LED illumination source and motorized stage. Time-lapse was carried out using cells seeded on Ibidi 8-well slides, and imaged at 37°C in L-15 medium/ 10% FBS using a 40X NA1.3 OIL objective. Epifluorescent stacks of fixed cells after processing by IF were acquired using 60X NA 1.0 OIL objective with 200 nm step. Image acquisition was controlled by Micro-Manager^54^ and images exported as tiff files.

### Image analysis and quantifications

Images were analysed using FIJI^55^, measuring net green intensity (T_i_) of cell after background subtraction at T0 and T200 mins. Picked cells which remained in prometaphase for the duration of the 200 mins. % degradation measured as (T0_I_ – T200_I_)/ T0_I._

Linescans were carried out using the BAR package in FIJI.

Mitochondrial lengths were analysed using MicroP^56^.

### AURKA and AURKB biochemical assays

AURKA and AURKB biochemical assays were performed as part of the ThermoFisher SelectScreen™ kinase profiling service (see full experimental protocol in Supplementary Information). Compounds were tested in AURKA or AURKB Z’-LYTE kinase assays using full-length purified protein (3 nM for AURKA, 20 nM for AURKB) and ATP at Km (10 μM for AURKA, 75 μM for AURKB). Across the three replicate assays, the assay Z’ averaged 0.87 for AURKA and 0.84 for AURKB.

### Statistics and Reproducibility

The number of biological repeats (two, three or four) for all experiments are indicated in figure legends. For live cell experiments data are presented either as mean values ±SDs from biological repeats, or pooled raw data as indicated in figure legends. For immunofluorescence analyses, two or three cover-slips were prepared for each condition as technical repeats, wth data pooled from technical repeats but not biological repeats. Quantified data analyses were plotted using GraphPad 6.01 (San Diego, CA, USA). Results were analyzed with ANOVA, Student’s *t*-test or Mann Whitney U test (non-parametric) as indicated in figure legends. Significant results are indicated as p < 0.05 (*), p ≤ 0.01 (**), p ≤ 0.001(***), p ≤ 0.0001 (****). Error bar values are stated as the mean ± SDs.

## Data availability

The authors declare that all data supporting the findings of this study are available within the paper and its Supplementary Information files. 1H and 13C NMR and HRMS analyses of the compounds described in this study are shown in the Supplementary Methods file. Quantified data are assembled are available in Supplementary Data 1 file.

## Acknowledgements

We thank Fanni Gergely and Laurent Pelletier for antibodies, Ian Storer, Zhongqian Sun and Zhengxu Li for Cpd A and D resynthesis and for design and synthesis of Cpd DX. Andreas Hock made valuable comments on the manuscript. RKW was supported by BBSRC-DTP, CA by a AstraZeneca-funded studentship and AMA by a Yousef Jameel Scholarship from the Cambridge International Trust. Work in CL’s lab is funded by BBSRC (BB/R004137/1).

## Author contributions

Study conceived and designed by CL and KR. TR synthesized compounds used. Experimental work was carried out and analysed by RKW, CA, AMA, AF. Manuscript written by CL, RKW, CA and revised by KR, IM, RKW, CA.

