## Supplementary Material for "Selective targeting of non-centrosomal AURKA functions through use of a targeted protein degradation tool"

**Supplementary Figures S1 – S7**
**Supplementary Methods (Chemical synthesis and validation of compounds)**

### Supplementary Figures

#### Figure S1

MLN8237

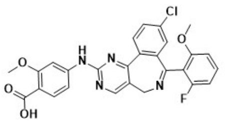

PROTAC-DX

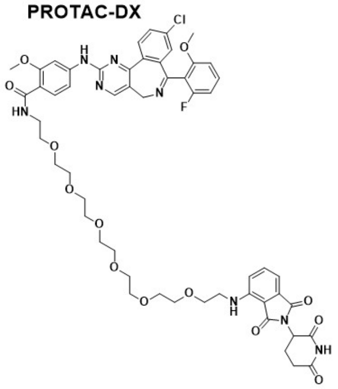

PROTAC-D

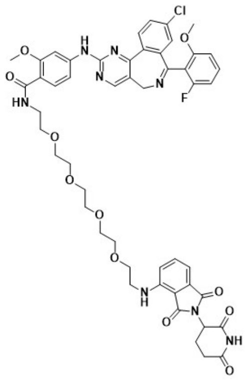

Cpd-A

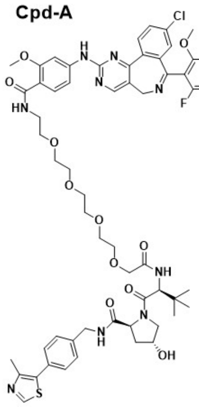

Chemical structures of compounds used in this study

#### Figure S2

| Cpd | AURKA $IC_{50}$ curve | AURKB $IC_{50}$ curve |
|---|---|---|
| MLN (alisertib) | 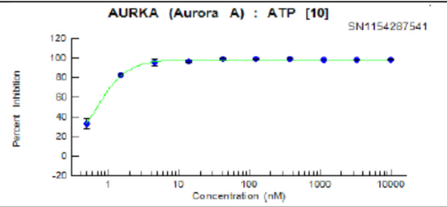 | 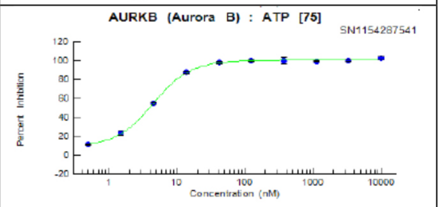 |
| PROTAC-D | 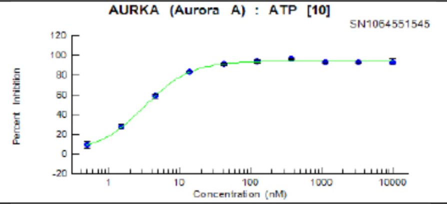 | 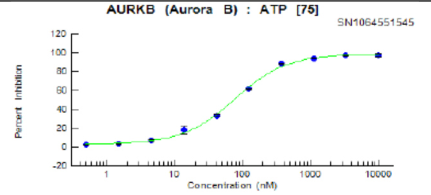 |
| PROTAC-DX | 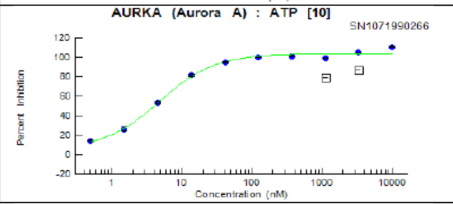 | 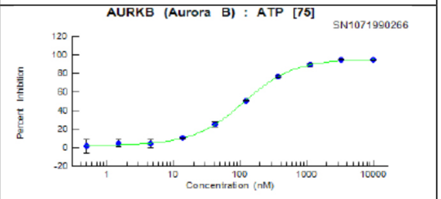 |
| CpdA | 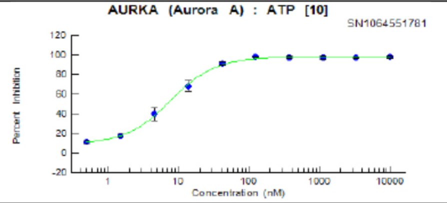 | 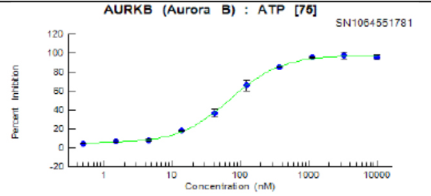 |

Biochemical assays of AURKA and AURKB inhibition by compounds used in this study. Representative dose-response curves are shown for each compound (n=3, except for CpdA, where n=2). $IC_{50}$s are reported in Table 2.

#### Figure S3

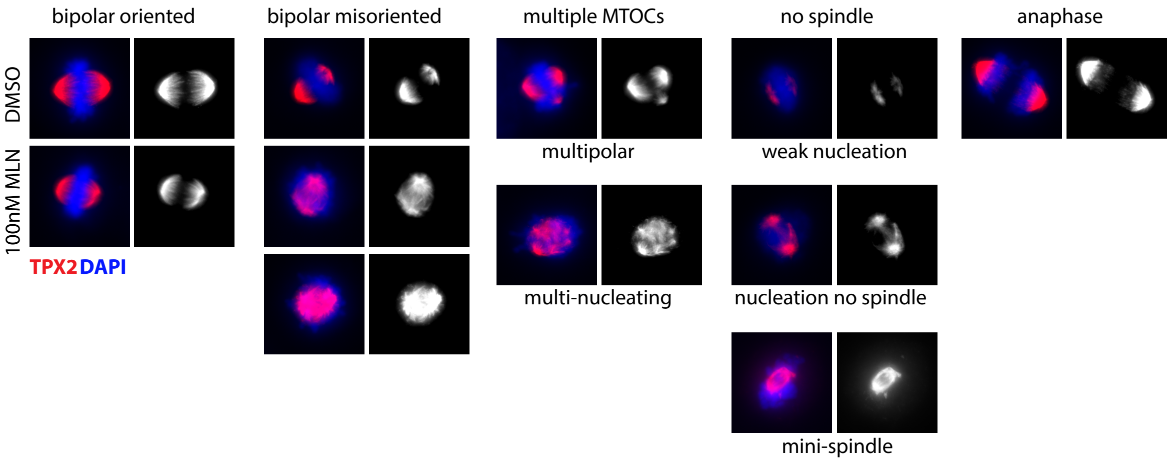


Examples of mitotic U2OS cells analysed in Figure 4D,E to illustrate different scoring categories applied. Cells were fixed 10 hours after release from single thymidine block and 3 hours after compound treatment, and stained for TPX2 (red), AURKA (green, not shown here) and DAPI (blue).

#### Figure S4

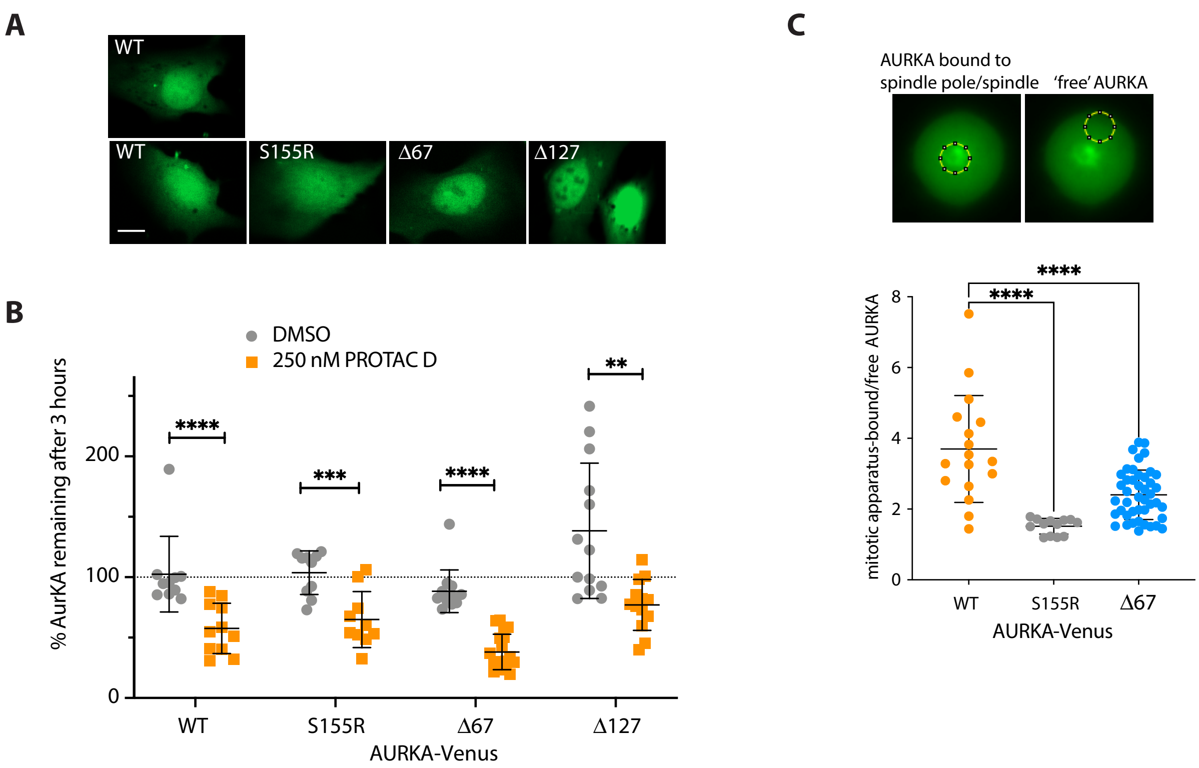


Response of exogenous AURKA-Venus to PROTAC-D treatment in U2OS cells transiently transfected with different versions of AURKA-Venus and treated for 16 hours with RO-3306 to arrest them in G2 phase. Cells were treated with 250nM PROTAC-D or DMSO as vehicle control and imaged by fluorescence timelapse microscopy over 3 hours. **A** Panels showing example cells transfected with each construct. **B** Degradation plots were derived by measuring AURKA-Venus levels in whole cells at the start and end of each experiment and calculating precentage protein remaining for each cell. Scatter plots show data from individual cells in a single experiment, with mean and SDs represented by error bars, and are representative of three repeats of the same experiment. The data show S155R sensitivity to PROTAC-D to be similar to that of wild-type protein, whilst Δ67 version appears more sensitive. Δ127 version accumulates more strongly than other versions in DMSO and is correspondingly less depleted by PROTAC-D treatment, presumably because of faulty proteostasis. ****, $p \leq 0.0001$ (Student's t-test, $n \geq 10$). **C** Quantification of AURKA-Venus localization at the mitotic apparatus (spindle or spindle pole of STLC-treated cells versus 'free' AURKA). Average pixel intensity in a circular ROI in the centre of the cell was divided by pixel values at the edge of the cell (top panels). Results from the cells analysed in Figure 6B are shown as scatter plots with means ± SDs indicated. ****, $p \leq 0.0001$, ordinary one-way ANOVA with Dunnett's post-hoc test.

#### Figure S5

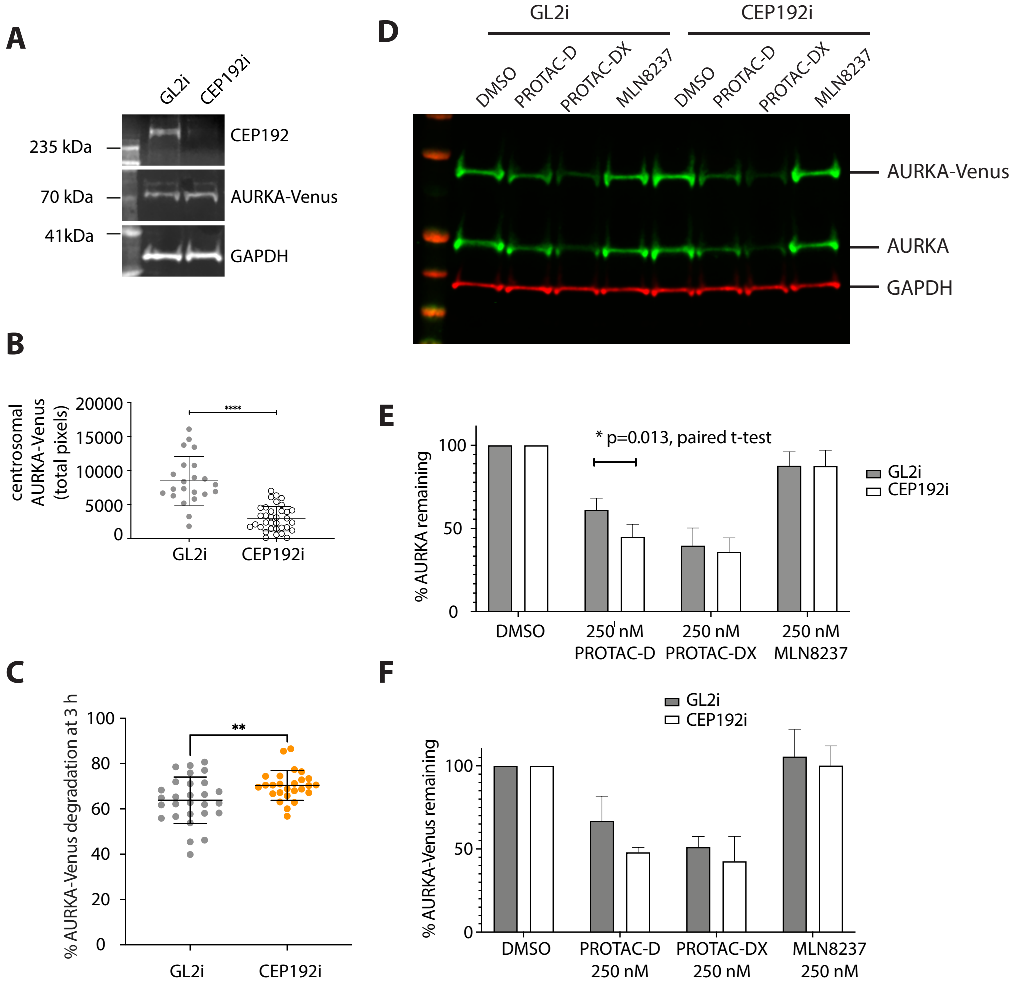


RPE1-AURKA-VenusKI cells were transfected for 48 hours with siRNA against CEP192 (CEP192i) or control siRNA (GL2i) then arrested in G2 phase with RO3306 for 16 hours before treatment with PROTAC-DX for 3 hours. Successful depletion of CEP192 was shown by immunoblot of cell extracts (**A**) and depletion of AURKA-VenusKI from centrosomes (as shown in Figure 6D) confirmed by measurement of average pixel intensity over a fixed region of interest around each centrosome, shown as scatter plot of values from individual centrosomes with mean ± SDs indicated (**B**). ****, p ≤ 0.0001 by unpaired t-Test for significance. **C** AurKA degradation in individual cells after 3 hours of treatment with 250 nM PROTAC-DX under conditions of CEP192i or GL2i is shown as scatter plots with mean ± SDs indicated. ** , p ≤ 0.01 by unpaired t-Test for significance, n ≥ 26. **D** Immunoblot of whole cell extracts also showed that depletion of AURKA and AURKA-Venus in response to PROTAC-D is enhanced by CEP192i, with the effect being quantitatively significant for endogenous AURKA (**E**) but not for AURKA-Venus (***F***). **A-D** show data from a single experiment and are representative of three independent experiments whilst **E,F** show mean ± SDs from quantified immunoblots of all 3 experiments (paired t-Test for significance).

#### Figure S6

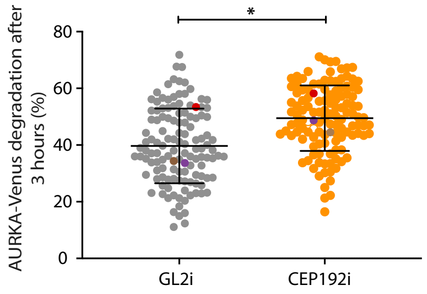


Graph accompanies that in Figure 6G, showing percentage degradation of AURKA-Venus in individual STLC-arrested cells after 3 hours treatment. Scatter plots contain pooled data from 3 independent experiments with mean values from each experiment overlaid in different colours. The third repeat of the experiment showed the same trend but with response to PROTAC reduced in all conditions (means shown in red). *, p ≤ 0.05, unpaired t-Test. n ≥ 124.

**Figure S7** Uncropped Li-COR imager scans of immunoblots used in Figures 1 - 3

Figure 1e

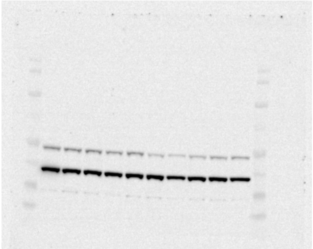

Figure 1g

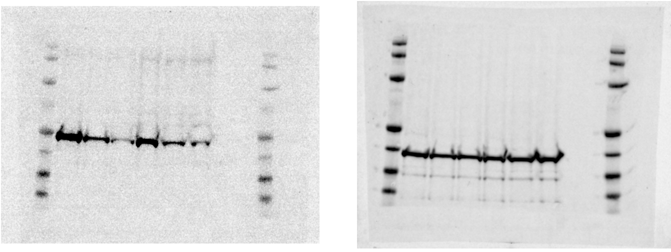

Figure 1i

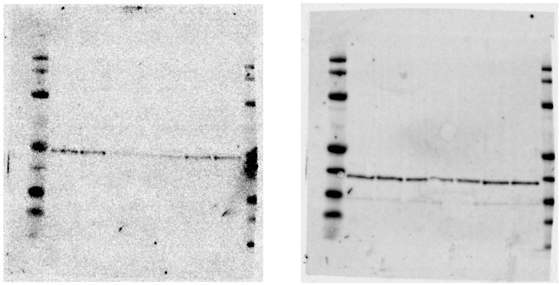

Figure 2c

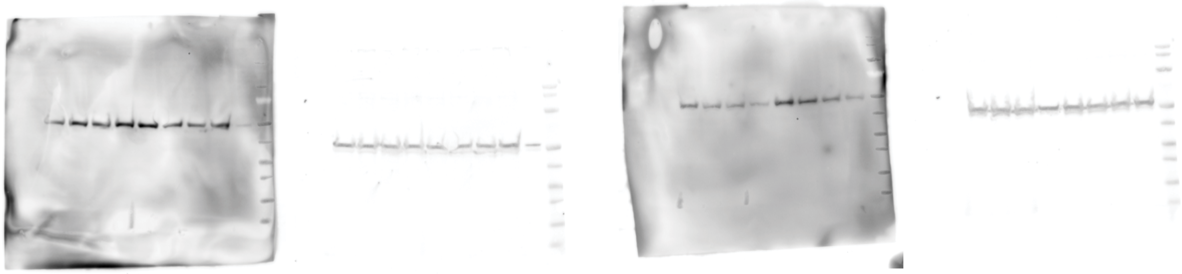

Figure 3b

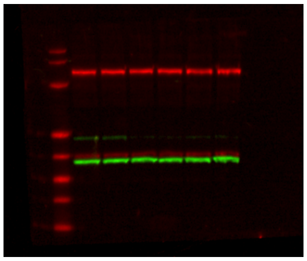

Figure 3d

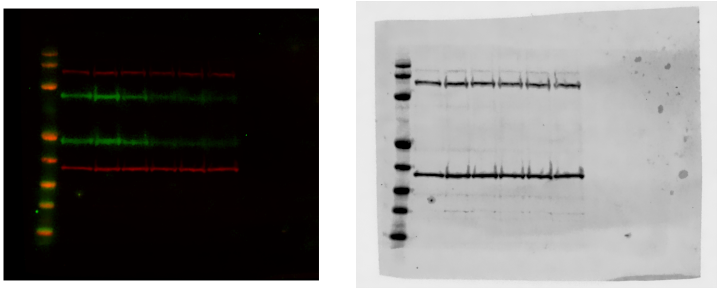

#### Supplementary Methods

##### Chemistry

***General chemistry experimental procedures***
Reagents and solvents (all anhydrous HPLC-grade) were obtained from commercial suppliers and used without any further purification unless otherwise stated. All reagents were weighed and handled in air unless otherwise stated. Brine refers to a saturated solution of NaCl. Concentration under reduced pressure refers to the use of a rotary evaporator.

***Spectroscopy***
$^{1}$H NMR and $^{13}$C spectra were recorded using Bruker Avance III, Avance III HD or Avance III NEO spectrometers at a proton frequency of 300 MHz, or a Bruker Avance Neo spectrometer at a proton frequency of 400 MHz, and are quoted in ppm for measurement against TMS or residual solvent peaks as internal standards. Unless otherwise stated, all experiments were carried out using DMSO-d6 as solvent. $^{1}$H NMR chemical shifts ($\delta$) are given in ppm ± 0.01, and coupling constants ($J$) are given in Hz ± 0.1 Hz. The $^{1}$H NMR spectra are reported as follows: $\delta$/ppm (multiplicity, coupling constant(s) $J$/Hz, number of protons). Multiplicity is abbreviated as follows: s = singlet, br s = broad singlet, d = doublet, br d = broad doublet, dd = doublet of doublets, t = triplet, dt = doublet of triplet, q = quartet, dq = doublet of quartet, quint = quintet, m = multiplet.

MS experiments were performed using a Waters Acquity system combined with a Waters Xevo Q-ToF Mass or a Shimadzu 2010EV UPLC system in ESI mode. LC was run in two setups: 1) BEH C18 column (1.7 µm, 2.1 x 50 mm) in combination with a gradient (2-95% B in 5 minutes) of aqueous 46 mM ammonium carbonate/ammonia buffer at pH 10 (A) and MeCN (B) at a flow rate of 1.0 mL/min or in combination with a gradient (5-95% B in 2 minutes) of water and TFA (0.05%) (A) and MeCN and TFA (0.05%) at a flow rate of 1.0 mL/min (B).

***Compound naming***
Compound names are those generated by ChemDraw 19.0.

***Compound purity***
All compounds detailed below were found to be > 93% pure as assessed by LCMS. All HPLC was performed on an Agilent 1200 system (Agilent, Santa Clara, CA) comprising 2 G1312B ultra-high-pressure binary pumps, a G1315C Diode Array detector, a G1316B Column Compartment, a G1379B Micro De-gasser, a G1367C Micro Well Plate Auto-sampler and a 35900E analog-to-digital converter. The eluent from the HPLC was split between an Agilent 6140 single quad mass spectrometer equipped with a multimode source and an ESA (ESA, Chelmsford, MA) Corona charged aerosol detector. HPLC reversed-phase separations were performed on a 50 × 2-mm Kinetex 2.6-µm C18 column (Phenomenex, Torrance, CA) at a flow rate of 700 µL per minute using a gradient comprising (A) HPLC grade water (VWR) with 0.05% formic acid (Sigma Aldrich) and (B) HPLC grade acetonitrile (Honeywell) with 0.05% formic acid. The conditions for pump 1 were at time 0, A = 90%, with a linear gradient such that after 2 min, B = 100%, which was held for 0.5 min before returning to starting conditions. The gradient conditions for pump 2 were the opposite of pump 1 so as to combine the 2 solvent streams after the column and produce a constant 50:50 volume/volume mix of A and B at the detectors. The Agilent 6140 MS acquired from 100 to 1000 Da in sequential positive and negative ion modes with a total cycle time of 1 s, and the output from the Corona CAD (ESA) detector was acquired at a rate of 5 Hz through the analog-to-digital converter. The diode array detector (DAD) scanned from 220 to 300 nm at a rate of 20 Hz. All data were acquired into Chemstation Version B.03.01 (Agilent). The data from the DAD and CAD detectors were integrated automatically within Chemstation.

| **Compound** | **Purity (%)** |
|---|---|
| **MLN8237 (alisertib)** | 100 |
| **D** | 100; 93.6 |
| **DX** | 98 |
| **A** | 100 |

**MLN8237 (alisertib)** was purchased from commercial vendors.

***Compound synthesis:***

*Synthesis of 4-((9-chloro-7-(2-fluoro-6-methoxyphenyl)-5H-benzo[c]pyrimido[4,5-e]azepin-2-yl)amino)-N-(14-((2-(2,6-dioxopiperidin-3-yl)-1,3-dioxoisoindolin-4-yl)amino)-3,6,9,12-tetraoxatetradecyl)-2-methoxybenzamide **(compound D)**:*

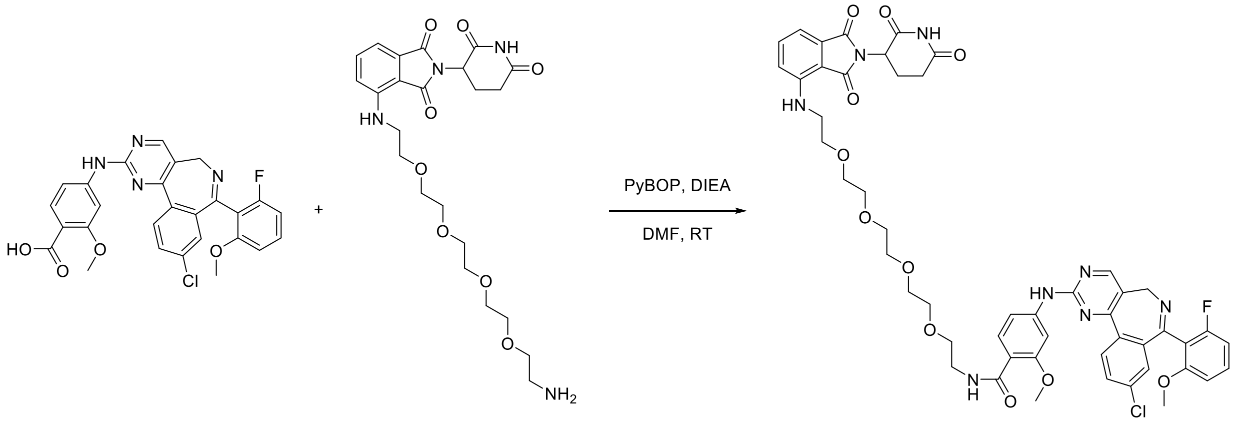


DIEA (20.0 µL, 0.115 mmol) was added to a mixture of 4-((14-amino-3,6,9,12-tetraoxatetradecyl)amino)-2-(2,6-dioxopiperidin-3-yl)isoindoline-1,3-dione hydrochloride[1,a] (20.0 mg, 0.0378 mmol), 4-((9-chloro-7-(2-fluoro-6-methoxyphenyl)-5*H*-benzo[*c*]pyrimido[4,5-*e*]azepin-2-yl)amino)-2-methoxybenzoic acid[b] (19.6 mg, 0.0378 mmol) and PyBOP (29.5 mg, 0.0567 mmol) in DMF (2 mL) at RT. The resulting mixture was stirred at RT for 4 hours. The crude product was directly purified by C18-flash chromatography (elution gradient 5 to 50% MeCN in water). Fractions containing the title compound were evaporated to dryness and repurified by preparative HPLC (Sunfire prep C18 column, 5µm, 30*150) using decreasingly polar mixtures of water (containing 0.05% TFA) and MeCN as eluents (elution gradient 45 to 55%) to afford 4-((9-chloro-7-(2-fluoro-6-methoxyphenyl)-5*H*-benzo[*c*]pyrimido[4,5-*e*]azepin-2-yl)amino)-*N*-(14-((2-(2,6-dioxopiperidin-3-yl)-1,3-dioxoisoindolin-4-yl)amino)-3,6,9,12-tetraoxatetradecyl)-2-methoxybenzamide (22.5 mg, 60%) as a yellow solid.

**$^{1}$H NMR** (400 MHz, DMSO-d6) δ 1.95 – 2.09 (m, 1H), 2.44 – 2.67 (m, 2H), 2.77 – 2.98 (m, 1H), 3.20 – 3.64 (m, 23H), 3.76 (br s, 1H), 3.93 (s, 3H), 4.86 (br s, 1H), 5.05 (dd, *J* = 12.8, 5.2 Hz, 1H), 6.57 (t, *J* = 5.3 Hz, 1H), 6.87 (br s, 2H), 7.01 (d, *J* = 7.0 Hz, 1H), 7.08 (d, *J* = 8.6 Hz, 1H), 7.22 (s, 1H), 7.35 – 7.46 (m, 2H), 7.53 (t, *J* = 7.8 Hz, 1H), 7.80 (d, *J* = 8.4 Hz, 1H), 7.87 (d, *J* = 8.6 Hz, 1H), 7.98 (s, 1H), 8.16 (t, *J* = 4.8 Hz, 1H), 8.29 (d, *J* = 8.5 Hz, 1H), 8.70 (s, 1H), 10.18 (s, 1H), 11.10 (s, 1H)

**$^{13}$C NMR**[c] (100 MHz, DMSO-d6) δ 22.2, 31.0, 41.7, 48.6, 49.5, 55.7, 56.3, 68.8, 69.1, 69.7, 69.76, 69.83, 101.4, 108.0, 109.2, 110.5, 110.6, 114.3, 117.3, 117.9 (d, $J_{CF}$ = 18.0 Hz), 123.4, 127.4, 130.2, 130.8, 131.1 (d, $J_{CF}$ = 9.9 Hz), 131.6, 132.0, 134.6, 135.2, 136.1, 137.6, 144.6, 146.4, 157.1, 157.7 (d, $J_{CF}$ = 6.8 Hz), 157.9, 159.4, 159.5 ($J_{CF}$ = 244.9 Hz), 160.2 (d, $J_{CF}$ = 1.7 Hz), 160.9, 164.2, 167.3, 168.9, 170.0, 172.7

**$^{19}$F NMR** (376 MHz, DMSO-d6) δ -117.2

**m/z** (ES+), [M+H]$^+$ = 993.2

**HRMS** (ES+) = calculated for $[C_{50}H_{51}ClFN_8O_{11}]^+$ [M+H]$^+$ = 993.3344, found = 993.3394

[a] Commercially available from specialist vendors

[b] Purchased from commercial vendors

[c] A number of peaks in the $^{13}$C spectrum overlay either with each other or with the solvent peak. Furthermore, some of the multiplets of the C-F coupled peaks are close to the baseline of the spectrum and thus are challenging to identify

*Synthesis of tert-butyl (20-((2-(2,6-dioxopiperidin-3-yl)-1,3-dioxoisoindolin-4-yl)amino)-3,6,9,12,15,18-hexaoxaicosyl)carbamate:*

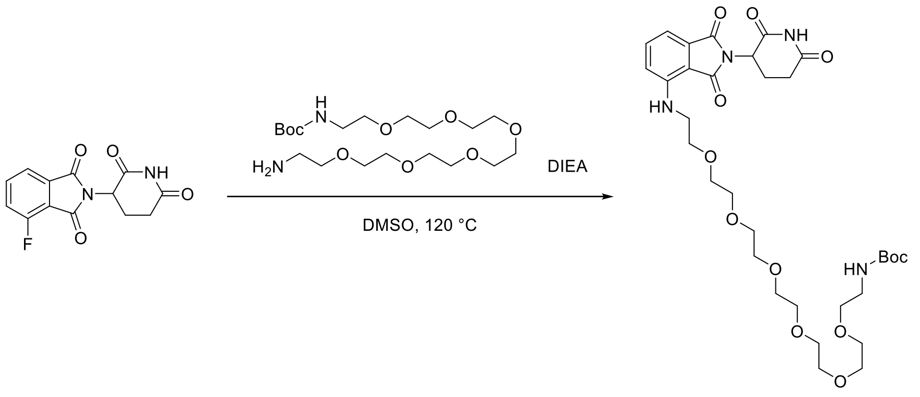


DIEA (1.52 mL, 8.70 mmol) was added to a solution of 2-(2,6-dioxopiperidin-3-yl)-4-fluoroisoindoline-1,3-dione (1.20 g, 4.34 mmol) and *tert*-butyl (20-amino-3,6,9,12,15,18-hexaoxaicosyl)carbamate (2.21 g, 5.21 mmol) in DMSO (30 mL) at RT. The resulting mixture was stirred at 120 °C for 16 hours. The reaction mixture was poured into water (100 mL) and extracted with EtOAc (2 x 100 mL). The organic layer was dried over anhydrous $Na_2SO_4$, filtered and concentrated under reduced pressure. The crude product was purified by flash column chromatography (elution gradient 0 to 100% EtOAc in petroleum ether). Pure fractions were evaporated to dryness to afford *tert*-butyl (20-((2-(2,6-dioxopiperidin-3-yl)-1,3-dioxoisoindolin-4-yl)amino)-3,6,9,12,15,18-hexaoxaicosyl)carbamate (2.30 g, 78%) as a yellow oil.

**$^1$H NMR** (300 MHz, Chloroform-d) δ 1.43 (s, 9H), 2.06 – 2.19 (m, 1H), 2.64 – 2.92 (m, 3H), 3.29 (br s, 2H), 3.46 (t, *J* = 5.3 Hz, 2H), 3.53 (t, *J* = 5.1 Hz, 2H), 3.57 – 3.68 (m, 20H), 3.72 (t, *J* = 5.4 Hz, 2H), 4.90 (dd, *J* = 12.1, 5.5 Hz, 1H), 5.07 (br s, 1H), 6.49 (br s, 1H), 6.92 (d, *J* = 8.5 Hz, 1H), 7.10 (d, *J* = 6.9 Hz, 1H), 7.49 (dd, *J* = 8.5, 7.2 Hz, 1H), 8.37 (br s, 1H)

**m/z** (ES+), $[M+Na]^+$ = 703.3

*Synthesis of 4-((20-amino-3,6,9,12,15,18-hexaoxaicosyl)amino)-2-(2,6-dioxopiperidin-3-yl)isoindoline-1,3-dione hydrochloride:*

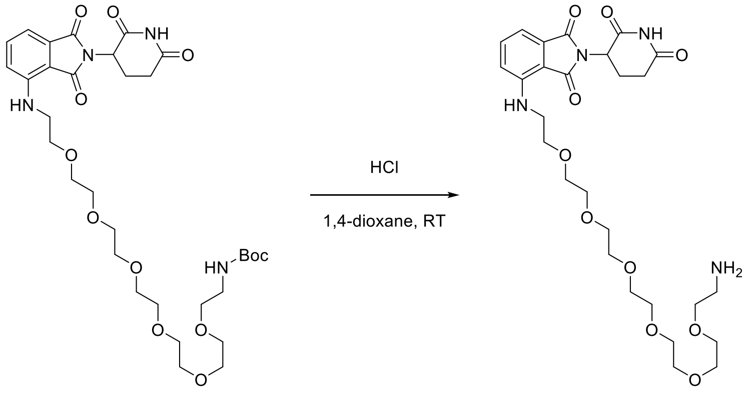


HCl in 1,4-dioxane (4M, 10.0 mL, 40.0 mmol) was added to a solution of *tert*-butyl (20-((2-(2,6-dioxopiperidin-3-yl)-1,3-dioxoisoindolin-4-yl)amino)-3,6,9,12,15,18-hexaoxaicosyl)carbamate (2.25 g, 3.31 mmol) in 1,4-dioxane (10 mL) at RT. The resulting mixture was stirred at RT for 4 hours. The solvent was removed under reduced pressure to afford 4-((20-amino-3,6,9,12,15,18-hexaoxaicosyl)amino)-2-(2,6-dioxopiperidin-3-yl)isoindoline-1,3-dione hydrochloride (1.72 g, 84%) as a yellow solid which was used in the subsequent step without any further purification.

**$^1$H NMR** (300 MHz, DMSO-d6) δ 1.95 – 2.11 (m, 1H), 2.52 – 2.65 (m, 2H), 2.79 – 3.02 (m, 3H), 3.42 – 3.68 (m, 26H), 5.06 (dd, *J* = 12.8, 5.4 Hz, 1H), 6.61 (br s, 1H), 7.04 (d, *J* = 7.0 Hz, 1H), 7.15 (d, *J* = 8.6 Hz, 1H), 7.59 (dd, *J* = 8.5, 7.1 Hz, 1H), 8.00 (s, 2H), 11.10 (s, 1H)

**m/z** (ES+), $[M+H]^+$ = 581.3

*Synthesis of 4-((9-chloro-7-(2-fluoro-6-methoxyphenyl)-5H-benzo[c]pyrimido[4,5-e]azepin-2-yl)amino)-N-(20-((2-(2,6-dioxopiperidin-3-yl)-1,3-dioxoisoindolin-4-yl)amino)-3,6,9,12,15,18-hexaoxaicosyl)-2-methoxybenzamide* ***(compound DX)****:*

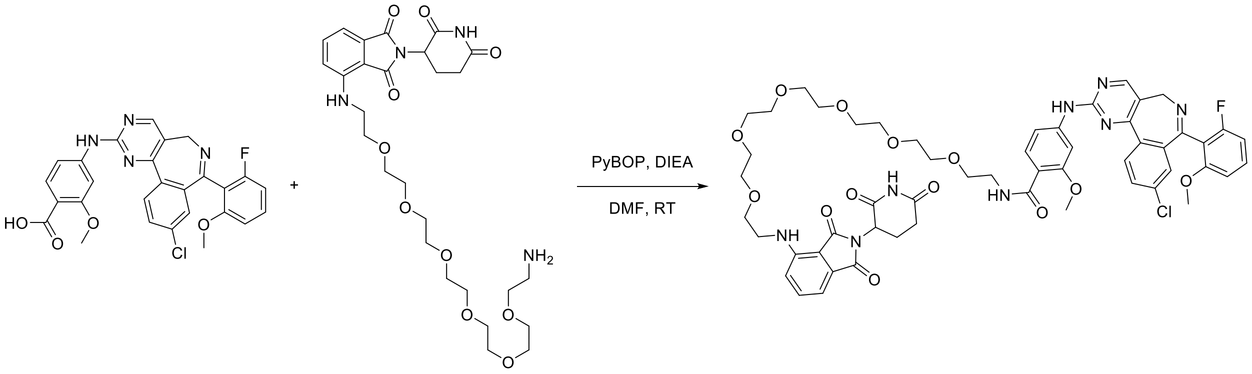


DIEA (20.0 µL, 0.115 mmol) was added to a mixture of 4-((20-amino-3,6,9,12,15,18-hexaoxaicosyl)amino)-2-(2,6-dioxopiperidin-3-yl)isoindoline-1,3-dione hydrochloride (23.8 mg, 0.0386 mmol), 4-((9-chloro-7-(2-fluoro-6-methoxyphenyl)-5*H*-benzo[*c*]pyrimido[4,5-*e*]azepin-2-yl)amino)-2-methoxybenzoic acid[d] (20.0 mg, 0.0385 mmol) and PyBOP (20.1 mg, 0.0386 mmol) in DMF (2 mL) at RT. The resulting mixture was stirred at RT for 6 hours. The crude product was directly purified by C18-flash chromatography (elution gradient 5 to 70% MeCN in water [containing 0.01% TFA]). Fractions containing the title compound were evaporated to dryness and repurified by preparative HPLC (Sunfire prep C18 column, 5 µm, 30*150) using decreasingly polar mixtures of water (containing 0.05% TFA) and MeCN (elution gradient 45 to 55%) to afford 4-((9-chloro-7-(2-fluoro-6-methoxyphenyl)-5*H*-benzo[*c*]pyrimido[4,5-*e*]azepin-2-yl)amino)-*N*-(20-((2-(2,6-dioxopiperidin-3-yl)-1,3-dioxoisoindolin-4-yl)amino)-3,6,9,12,15,18-hexaoxaicosyl)-2-methoxybenzamide (10.7 mg, 26%) as a yellow solid.

**$^1$H NMR** (400 MHz, DMSO-d6) δ 1.97 – 2.07 (m, 1H), 2.46 – 2.64 (m, 2H), 2.88 (ddd, *J* = 17.9, 14.1, 5.2 Hz, 1H), 3.01 – 3.66 (m, 31H), 3.79 (br s, 1H), 3.93 (s, 3H), 4.87 (br s, 1H), 5.05 (dd, *J* = 12.8, 5.2 Hz, 1H), 6.58 (t, *J* = 5.4 Hz, 1H), 6.88 (br s, 2H), 7.02 (d, *J* = 7.0 Hz, 1H), 7.10 (d, *J* = 8.6 Hz, 1H), 7.22 (s, 1H), 7.36 – 7.49 (m, 2H), 7.55 (t, *J* = 7.8 Hz, 1H), 7.80 (d, *J* = 8.4 Hz, 1H), 7.87 (d, *J* = 8.6 Hz, 1H), 7.98 (s, 1H), 8.16 (t, *J* = 4.8 Hz, 1H), 8.29 (d, *J* = 8.5 Hz, 1H), 8.70 (s, 1H), 10.18 (s, 1H), 11.09 (s, 1H)

**$^{13}$C NMR**[e] (100 MHz, DMSO-d6) 22.1, 31.0, 41.7, 48.6, 49.5, 55.7, 56.3, 68.9, 69.1, 69.7, 69.75, 69.80, 69.82, 101.4, 108.0, 109.2, 110.5, 110.6, 114.3, 117.4, 117.9 (d, $J_{CF}$ = 18.5 Hz), 123.4, 127.4, 130.2, 130.8, 131.1 (d, $J_{CF}$ = 10.0 Hz), 131.6, 132.1, 134.6, 135.1 (d, $J_{CF}$ = 2.8 Hz), 136.1, 137.7, 144.6, 146.4, 157.1, 157.7 (d, $J_{CF}$ = 6.7 Hz), 157.9, 159.4, 159.5 (d, $J_{CF}$ = 244.8 Hz), 160.2 (d, $J_{CF}$ = 2.2 Hz), 160.9, 164.2, 167.3, 168.9, 170.0, 172.7

**$^{19}$F NMR** (376 MHz, DMSO-d6) δ -117.2

**m/z** (ES+), $[M+H]^+$ = 1082.5

**HRMS** (ES+) = calculated for $[C_{54}H_{59}ClFN_8O_{13}]^+$ $[M+H]^+$ = 1081.3869, found = 1081.3926

---

[d] Purchased from commercial vendors

[e] A number of peaks in the $^{13}$C spectrum overlay either with each other or with the solvent peak. Furthermore, some of the multiplets of the C-F coupled peaks are close to the baseline of the spectrum and thus are challenging to identify

*Synthesis of (2S,4R)-1-((S)-18-(tert-butyl)-1-(4-((9-chloro-7-(2-fluoro-6-methoxyphenyl)-5H-benzo[c]pyrimido[4,5-e]azepin-2-yl)amino)-2-methoxyphenyl)-1,16-dioxo-5,8,11,14-tetraoxa-2,17-diazanonadecan-19-oyl)-4-hydroxy-N-(4-(4-methylthiazol-5-yl)benzyl)pyrrolidine-2-carboxamide (**compound A**)*

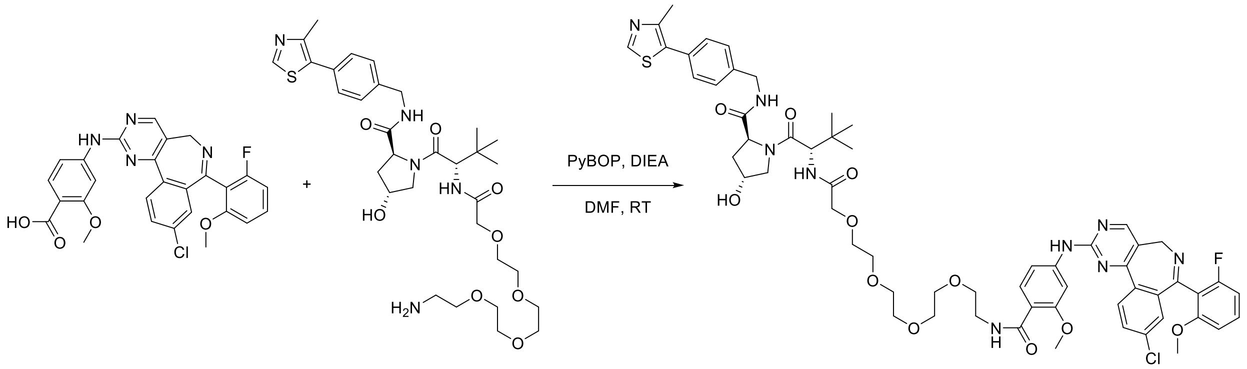


DIEA (135 µL, 0.773 mmol) was added to a mixture of (2*S*,4*R*)-1-((*S*)-17-amino-2-(*tert*-butyl)-4-oxo-6,9,12,15-tetraoxa-3-azaheptadecanoyl)-4-hydroxy-*N*-(4-(4-methylthiazol-5-yl)benzyl)pyrrolidine-2-carboxamide (128 mg, 0.193 mmol), 4-((9-chloro-7-(2-fluoro-6-methoxyphenyl)-5*H*-benzo[*c*]pyrimido[4,5-*e*]azepin-2-yl)amino)-2-methoxybenzoic acid[f] (100 mg, 0.193 mmol) and PyBOP (150 mg, 0.288 mmol) in DMF (5 mL) at RT. The resulting mixture was stirred at RT for 16 hours. The solvent was removed under reduced pressure and the crude product was purified by C18-flash chromatography (elution gradient 0 to 60% MeCN in water [containing 0.01% FA]). Fractions containing the title compound were evaporated to dryness and repurified by preparative HPLC (Sunfire prep C18 column, 5 µm, 30*150) using decreasingly polar mixtures of water (containing 0.1% FA) and MeCN (elution gradient 44 to 55%) to afford (2*S*,4*R*)-1-((*S*)-18-(*tert*-butyl)-1-(4-((9-chloro-7-(2-fluoro-6-methoxyphenyl)-5*H*-benzo[*c*]pyrimido[4,5-*e*]azepin-2-yl)amino)-2-methoxyphenyl)-1,16-dioxo-5,8,11,14-tetraoxa-2,17-diazanonadecan-19-oyl)-4-hydroxy-*N*-(4-(4-methylthiazol-5-yl)benzyl)pyrrolidine-2-carboxamide (90.0 mg, 40%) as a white solid.

**$^{1}$H NMR** (400 MHz, DMSO-d6) δ 0.94 (s, 9H), 1.86 – 1.96 (m, 1H), 2.02 – 2.11 (m, 1H), 2.43 (s, 3H), 3.20 – 3.74 (m, 22H), 3.92 (s, 3H), 3.96 (s, 2H), 4.25 (dd, *J* = 15.7, 5.5 Hz, 1H), 4.32 – 4.50 (m, 3H), 4.57 (d, *J* = 9.6 Hz, 1H), 4.84 (br s, 1H), 5.17 (br s, 1H), 6.86 (br s, 2H), 7.22 (s, 1H), 7.32 – 7.50 (m, 7H), 7.76 – 7.83 (m, 1H), 7.87 (d, *J* = 8.6 Hz, 1H), 7.97 (s, 1H), 8.16 (t, *J* = 4.8 Hz, 1H), 8.29 (d, *J* = 8.5 Hz, 1H), 8.59 (t, *J* = 5.7 Hz, 1H), 8.70 (s, 1H), 8.95 (s, 1H), 10.17 (s, 1H)

**$^{13}$C NMR**[g] (100 MHz, DMSO-d6) 15.9, 26.2, 35.7, 37.9, 41.7, 49.5, 55.7, 56.3, 56.6, 58.8, 68.9, 69.1, 69.6, 69.7, 69.83, 69.85, 69.87, 70.5, 101.5, 108.0, 110.5, 114.3, 117.9 ($J_{CF}$ = 17.4 Hz), 123.4, 127.5, 128.7, 129.7, 130.2, 130.9, 131.1, 131.2 ($J_{CF}$ = 11.1 Hz), 131.6, 134.6, 135.2, 137.7, 139.4, 144.7, 147.7, 151.4, 157.1, 157.8 ($J_{CF}$ = 7.1 Hz), 157.9, 159.4, 159.6 ($J_{CF}$ = 244.8 Hz), 160.2, 160.9, 164.3, 168.6, 169.2, 171.8

**$^{19}$F NMR** (376 MHz, DMSO-d6) δ -117.2

**m/z** (ES+), $[M+H]^+$ = 1164.4

**HRMS** (ES+) = calculated for $[C_{59}H_{68}ClFN_9O_{11}S]^+$ $[M+H]^+$ = 1164.4426, found = 1164.4482

---

[f] Purchased from commercial vendors

[g] A number of peaks in the $^{13}$C spectrum overlay either with each other or with the solvent peak. Furthermore, some of the multiplets of the C-F coupled peaks are close to the baseline of the spectrum and thus are challenging to identify

**NMR spectra for compound D:**

**[1]H NMR** (400 MHz, DMSO-d6)

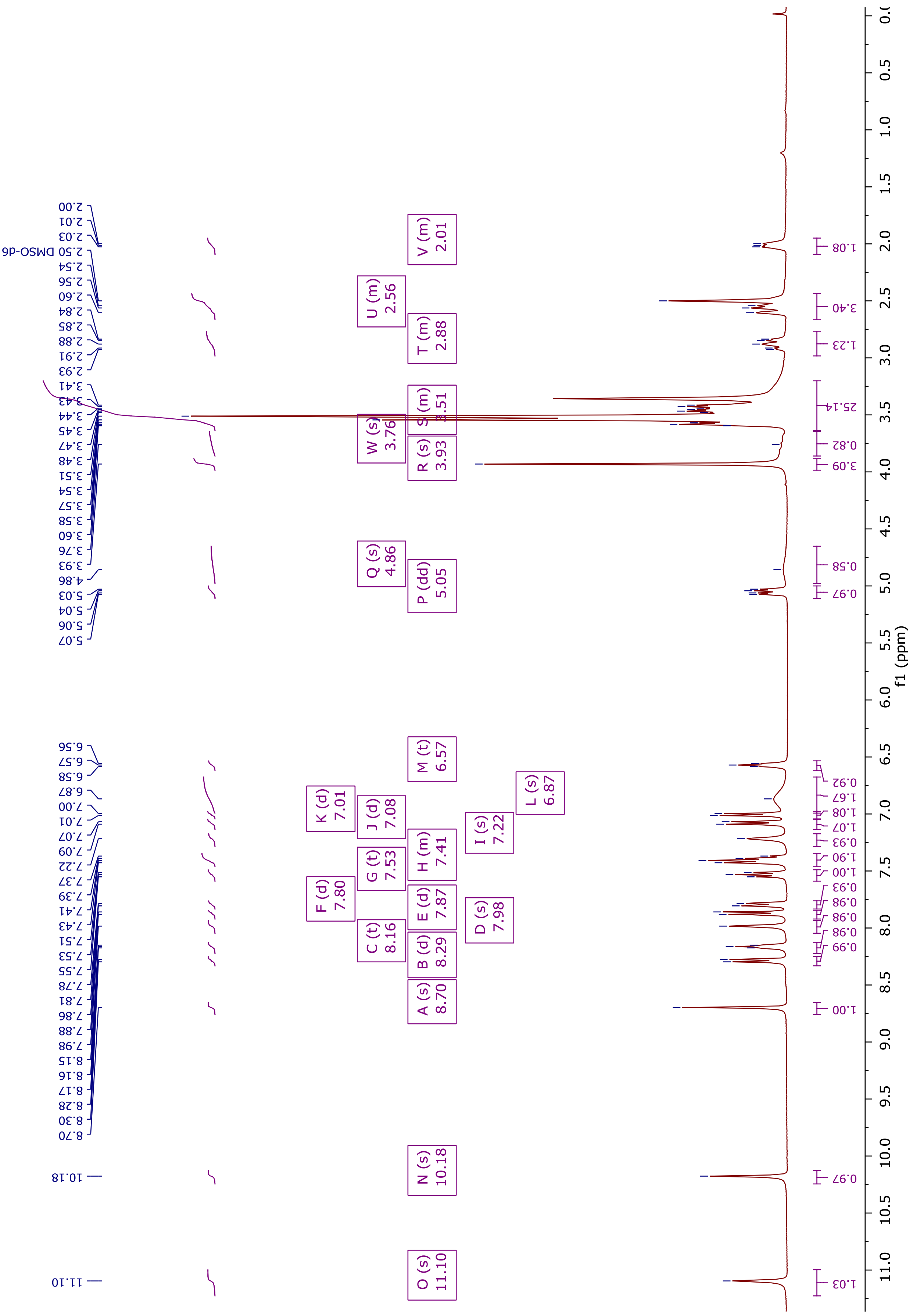

**$^{13}$C NMR** (100 MHz, DMSO-d6)[h]

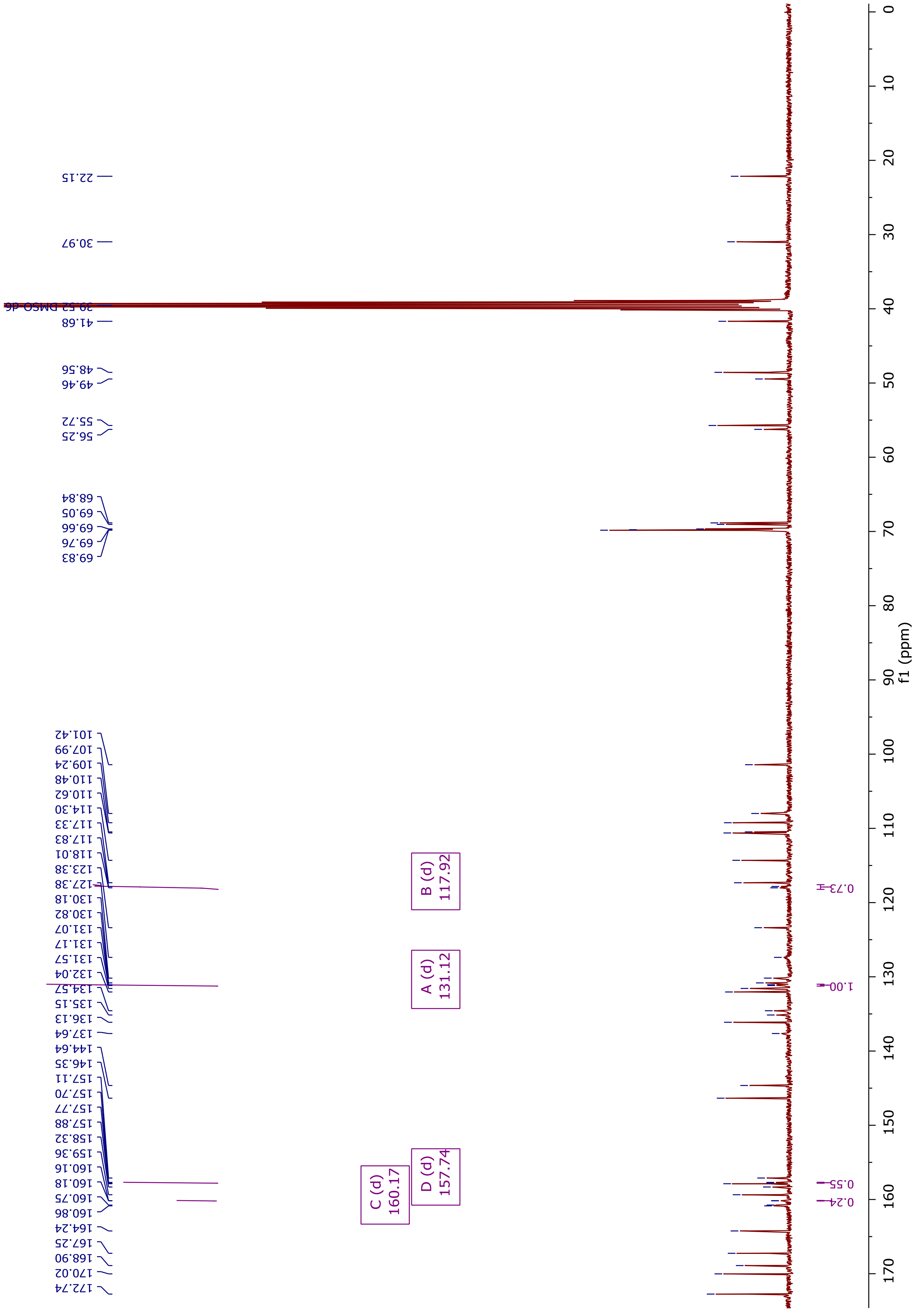


[h] The doublet 159.5 ($J_{CF}$ = 244.9 Hz) has been picked as two peaks for clarity

**NMR spectra for compound DX:**

$^1$**H NMR** (400 MHz, DMSO-d6)

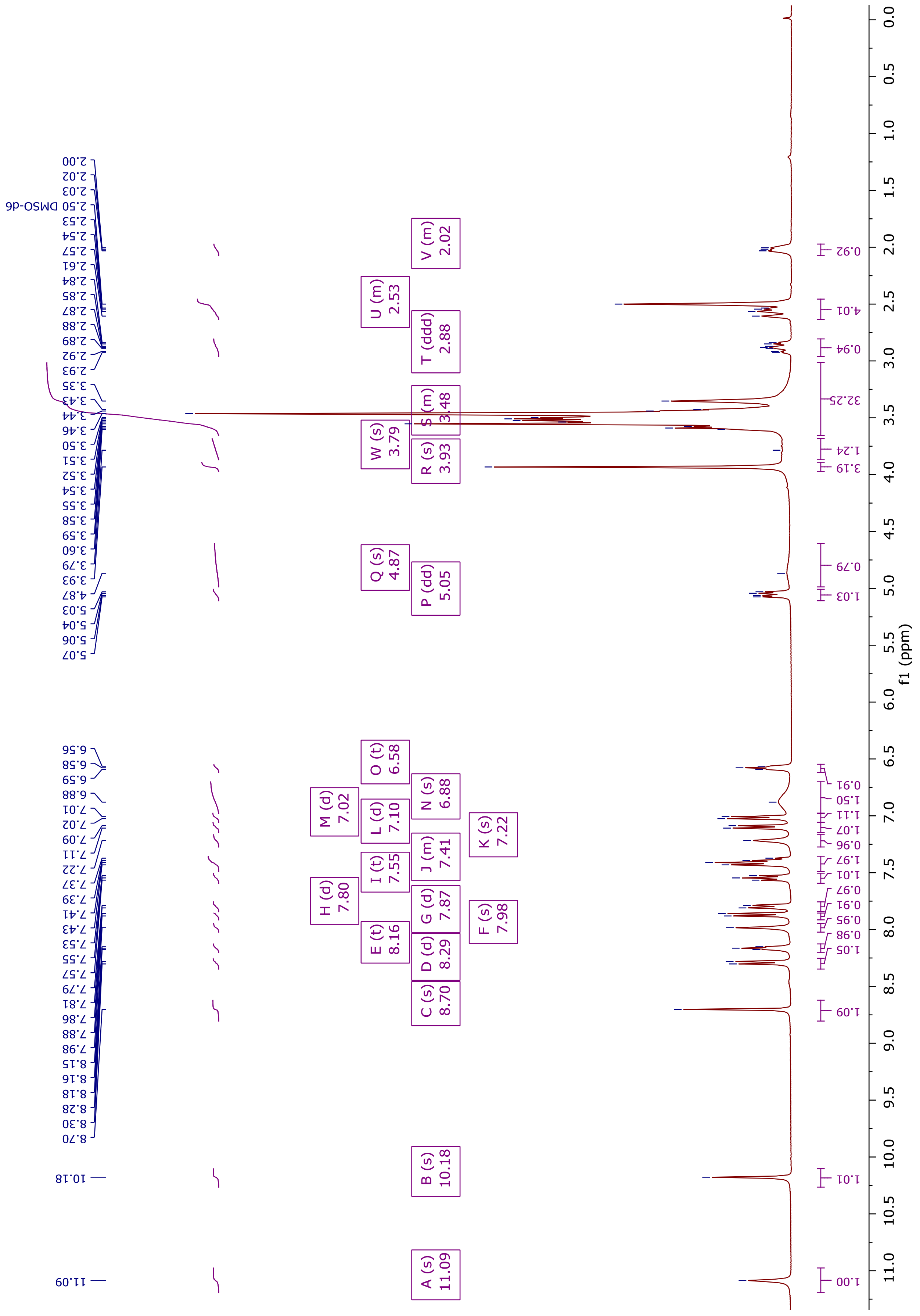

**$^{13}$C NMR** (100 MHz, DMSO-d6)[i]

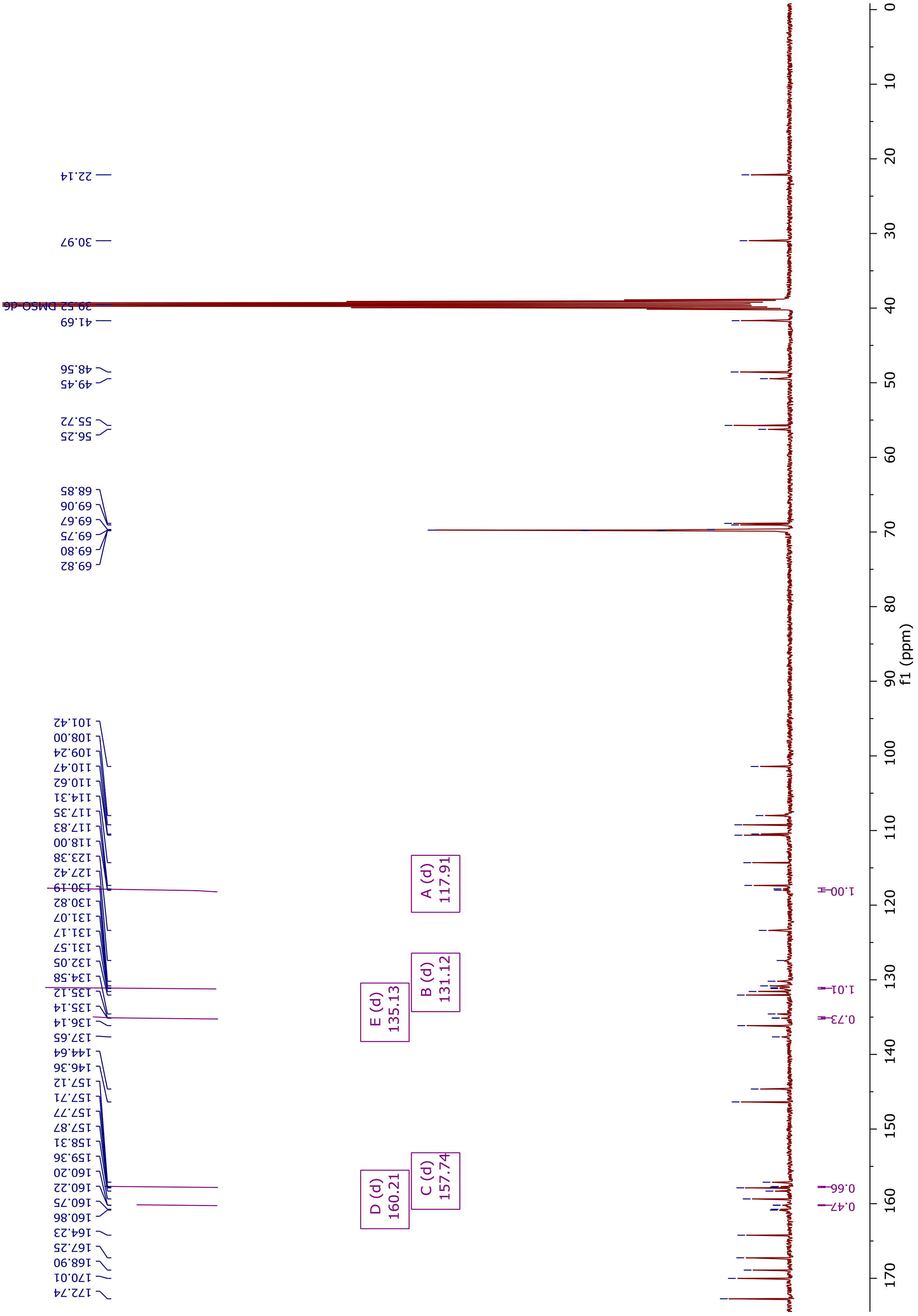


[i] The doublet 159.5 (d, $J_{CF}$ = 244.8 Hz) has been picked as two peaks for clarity

**NMR spectra for compound A:**

**$^{1}$H NMR** (400 MHz, DMSO-d6)

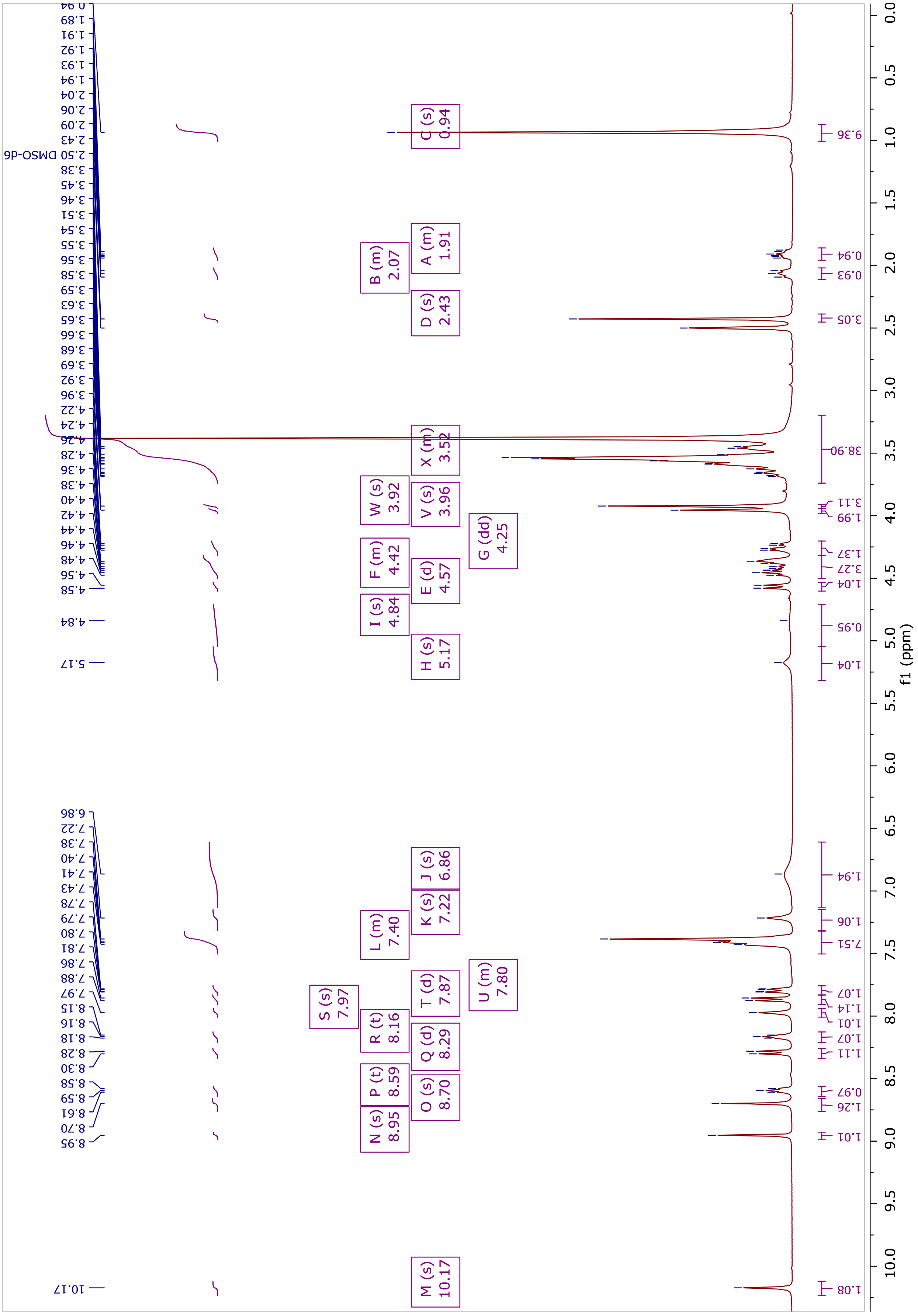

**$^{13}$C NMR** (100 MHz, DMSO-d6)[j]

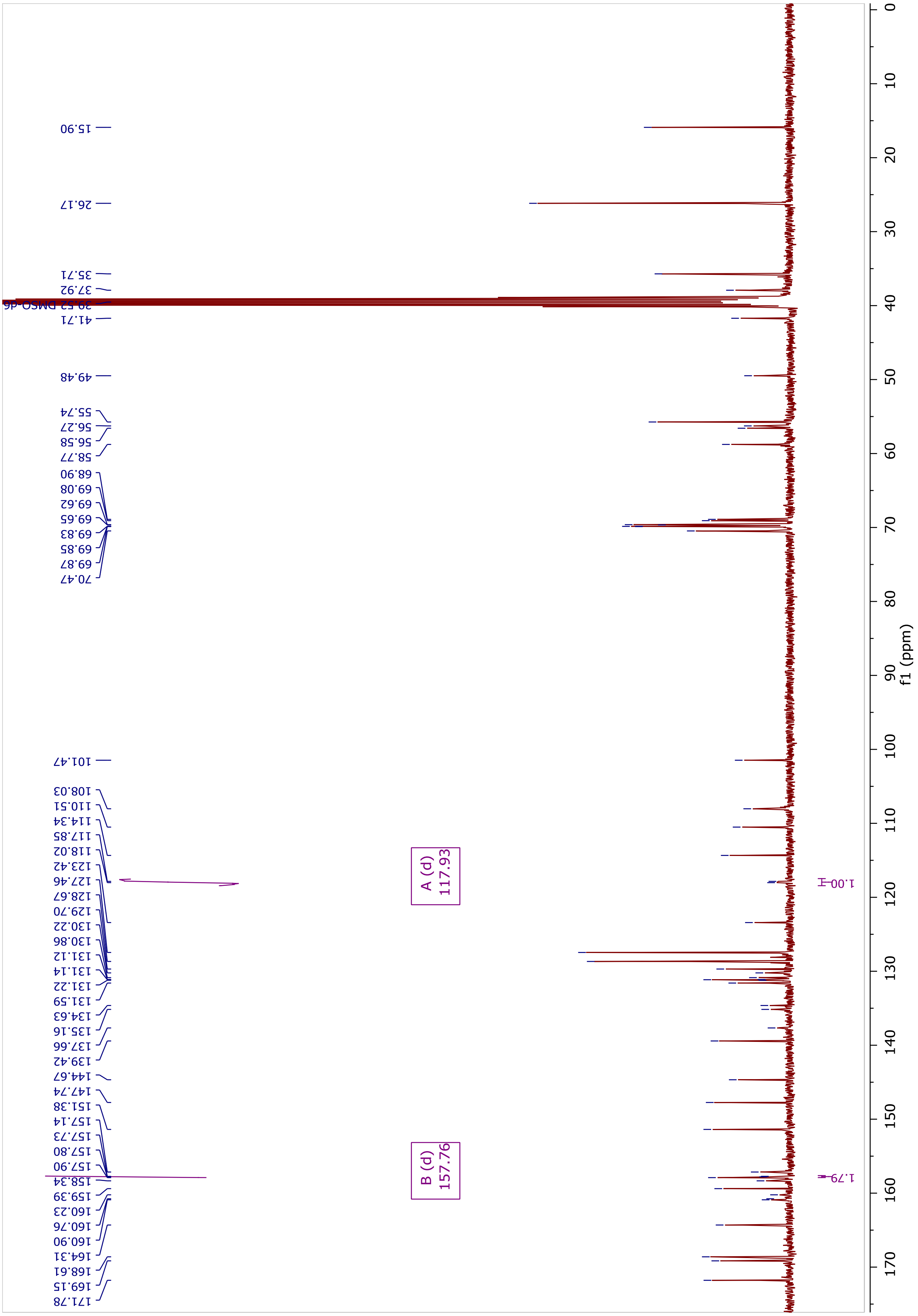


[j] The doublets 131.2 (d, $J_{CF}$ = 11.1 Hz) and 159.6 ($J_{CF}$ = 244.8 Hz) have both been picked as two peaks respectively for clarity

##### Chemistry abbreviations

| | |
|---|---|
| **°C** | Degrees Celsius |
| **Ac** | Acetyl |
| **App** | Apparent |
| **Aq** | Aqueous |
| **Ar** | Aryl |
| **Bn** | Benzyl |
| **Boc** | *tert-butoxycarbonyl* |
| **Bp** | Boiling point |
| **br** | Broad |
| **D** | Doublet |
| **dd** | Doublet of doublets |
| **DIEA** | *N*,*N*-Diisopropylethylamine/Hünig's base |
| **DMF** | Dimethylformamide |
| **DMSO** | Dimethyl sulfoxide |
| **Dppf** | [1,1′-Bis(diphenylphosphino)ferrocene] |
| **EDTA** | Ethylenediaminetetraacetic acid |
| **Equiv** | Equivalents |
| **$ES^-$** | Negative mode ESI |
| **$ES^+$** | Positive mode ESI |
| **ESI** | Electrospray ionization |
| **EtOAc** | Ethyl acetate |
| **FA** | Formic acid |
| ***J*** | Coupling constant |
| **M** | Molar concentration |
| **m** | Multiplet |
| **m/z** | Mass to charge ratio |
| **Me** | Methyl |
| **MeCN** | Acetonitrile |
| **MeOH** | Methanol |
| **mg** | Milligram |
| **MHz** | Mega hertz |
| **Min** | Minute(s) |
| **mL** | Millilitre |
| **mM** | millimolar |
| **mmol** | Millimole |
| **Mol** | mole |
| **MS** | Low resolution mass spectrometry |
| **NMR** | Nuclear Magnetic Resonance |
| **o** | Ortho |
| **p** | para |
| **PE** | Petroleum ether bp 60-90 °C |
| **Ph** | Phenyl |
| **ppm** | Parts per million |
| **PyBOP** | Benzotriazole-1-yl-oxy-tris-pyrrolidino-phosphonium hexafluorophosphate |
| **q** | Quartet |
| **quint** | Quintet |
| **rt** | Room temperature |
| **TFA** | Trifluoroacetic acid |
| **TMS** | Tetramethylsilane |
| **µL** | Microlitre |

**Supporting information references and notes**

1. Synthesised using reported conditions: (a) For HCl salt form: Mainolfi, N.; Ji, N.; Kluge, A. F.; Weiss, M. M.; Zhang, Y. Preparation of peptide conjugates as IRAK4 protein kinase degraders. US20190192668A1, 2019. (b) for TFA salt form: Peng, L.; Zhang, Z.; Lei, C.; Li, S.; Zhang, Z.; Ren, X.; Chang, Y.; Zhang, Y.; Xu, Y.; Ding, K., Identification of New Small-Molecule Inducers of Estrogen-related Receptor α (ERRα) Degradation. *ACS medicinal chemistry letters* **2019**, *10* (5), 767-772.
